# CD90 is not constitutively expressed in functional innate lymphoid cells

**DOI:** 10.1101/2021.12.11.472210

**Authors:** J-H Schroeder, G Beattie, JW Lo, T Zabinski, RG Jenner, N Powell, J F Neves, GM Lord

## Abstract

Huge progress has been made in understanding the biology of innate lymphoid cells (ILC) by adopting several well-known concepts in T cell biology. As such, flow cytometry gating strategies and markers, such as CD90, to identify ILC have been applied. Here, we report that most non-NK intestinal ILC have a high expression of CD90 as expected, but surprisingly a sub-population of cells exhibit low or even no expression of this marker. CD90-negative and CD90-low CD127^+^ ILC were present amongst all ILC subsets in the gut. The frequency of CD90-negative and CD90-low CD127^+^ ILC was dependent on stimulatory cues *in vitro* and enhanced due to dysbiosis *in vivo*. CD90-negative and CD90-low CD127^+^ ILC played a functional role as a source of IL-13, IFNγ and IL-17A at steady state and upon dysbiosis- and dextran sulphate sodium-elicited colitis. Hence, this study reveals that, contrary to expectations, CD90 is not constitutively expressed by functional ILC in the gut.

## INTRODUCTION

Resident leukocytes play an important role in maintaining mucosal surfaces at steady state and early during an infection (Bal *et al*., 2020, Klose *et al*., 2020). Since the discovery of innate lymphoid cells (ILC) about a decade ago, it has become increasingly apparent that these cells play a significant role in mucosal homeostasis. However, the role for ILCs is far from being fully characterised, and much of the current knowledge has been gained from testing concepts that had previously been established for T and NK cell biology. As such, group 1, 2 and 3 ILC (ILC1, ILC2 and ILC3) express T-bet, GATA3 and RORγt, respectively, as characteristic transcription factors as well as cytokines associated with Th1, Th2 and Th17 cells (Bal *et al*., 2020, Schroeder *et al*., 2022). Due to the absence of TCR expression in ILC, these cells elicit immune functions in response to cytokines, chemokines and neurotransmitters, as has been well described for NK cells (Bal *et al*., 2020, Klose *et al*., 2020).

Similarly to T and NK cells, ILC express the glycophosphatidylinositol (GPI) anchored protein CD90 in diverse tissues, and CD90 has often been used as a key marker to identify ILC (Sauzay *et al*., 2019, Leyton *et al*., 2019, Barman *et al*., 2021, Cox *et al*., 2021, Fachi *et al*., 2021, Ualiyeva *et al*., 2021, He *et al*., 2021, Xiao *et al*., 2022, He *et al*., 2022, Han *et al*., 2022, Peng *et al*., 2022, Chen *et al*., 2022, Wu *et al*., 2022a, Wu *et al*., 2022b, Glaubitz *et al*., 2022, Liu *et al*., 2022, Bruchard *et al*., 2022, Sheikh *et al*., 2022, Riding *et al*., 2022, Peng *et al*., 2022, Schmalzl *et al*., 2022), and CD90 has been used as key target to deplete ILC in Rag-deficient mice using a specific antibody (e.g. Powell *et al*., 2012, Mortha *et al*., 2014, Martin *et al*., 2017, Rafei-Shamsabadi *et al*., 2018, Castro-Dopico *et al*., 2020, Ray *et al*., 2022, Dobeš *et al*., 2022, Zhou *et al*., 2022). Despite the presence of CD90 on T and NK cells, very little is known regarding its functionality (Sauzay *et al*., 2019). In NK cells, CD90 downregulation was associated with successful differentiation, but its presence has also been linked to an activation phenotype (Gillard *et al*., 2011, Alvarez *et al*., 2019, Potempa *et al*., 2022). IL-17A-producing inflammatory ILC2 in lungs and small intestinal lamina propria (SI LP) have been observed to have lower expression of CD90 in comparison to natural ILC2, but the implications of this are not known (Huang *et al*., 2014, Flamar *et al*., 2020, Roberts *et al*., 2021). In relation to this, transition of CD90^low^ to CD90^high^ ILC2 precursors has been described using an *in vitro* model in which CLP were seeded, but again the role of the gain in CD90 is unknown (Seehus *et al*., 2014). Recently, it was reported that in the murine liver Ly49E^+^ ILC1 have a lower expression of CD90 than Ly49E^-^ ILC1 (Chen *et al*., 2022, Sparano *et al*., 2022).

Here, we report for the first time that intestinal lamina propria ILC exhibit varied expression of CD90, and strikingly some ILC show no expression of this marker. These CD90^-^ and CD90^low^ ILC are a significant source of IFNγ, IL-13 and IL-17A upon dysbiosis and dextran sulphate sodium (DSS)-elicited colitis. However, in naïve mice, CD90^-^ ILC have a dominant type 2 cytokine expression profile. Furthermore, stimulation with IL-25/IL-33 promotes the frequency of CD90^-/low^ ILC2 *in vitro*. Conversely, IL-12/IL-18 stimulation results in a lower prevalence of CD90^-/low^ NKp46^+^ ILC. These data suggest that CD90 expression in intestinal ILC is regulated by cytokines and has a limited suitability as a constitutive marker of the ILC lineage.

## RESULTS

### CD90-negative colonic lamina propria CD127^+^ ILC produce cytokines upon induced colitis

CD90 expression in ILC was tested in a mouse model of DSS-induced colitis. BALB/c *Rag2*^-/-^ mice were treated with 3% DSS in the drinking water for 5 days after which the animals showed clinical signs of colitis like weight loss (Schroeder *et al*., 2021a), and the cytokine expression profile of colonic lamina propria (cLP) ILC was analysed at day 10. Analyses of Lin^-^ CD45^+^ (CD3, CD5, CD45R, CD19, CD11b, TER-119, Gr-1, FcεRI) CD127^+^ colonic lamina propria (cLP) ILC re-stimulated with PMA and ionomycin (PI) *in vitro* revealed that in addition to CD90^high^ ILC there were CD90^-^ and CD90^low^ ILC populations (Figure 1a, Supplementary Figure 1a, b).

**Figure 1.**
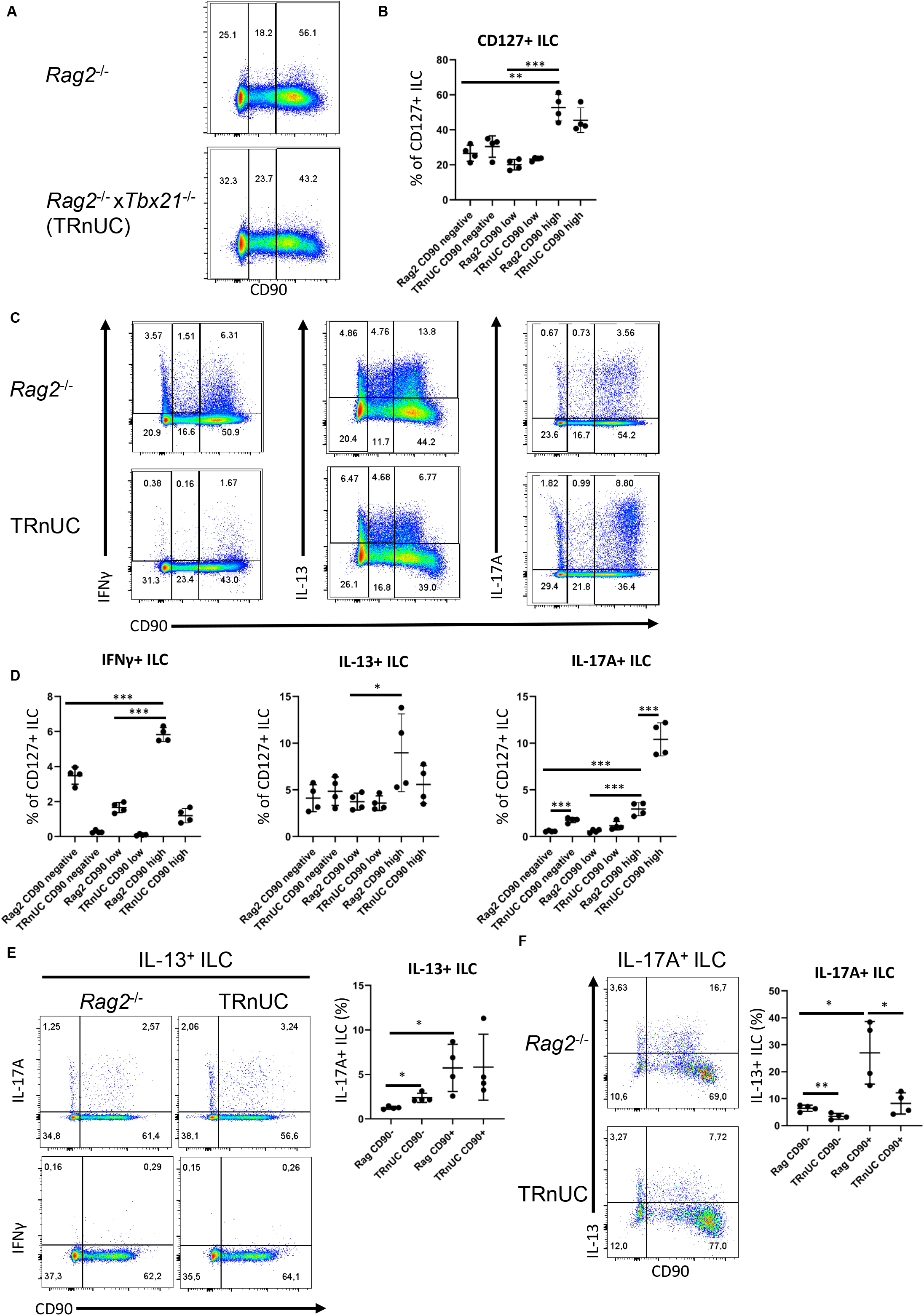
CD90-negative Rag-deficient ILC are a substantial source of IFNγ and IL-13 during DSS colitis. cLP ILC from 5% DSS-treated *Rag2*^-/-^ and TRnUC mice were isolated and stimulated with PMA and ionomycin (3 hours) prior to flow cytometry analysis. (a) Frequencies of CD90^hi^, CD90^low^ and CD90^-^ in total CD127^+^ ILC and (b) statistical analyses are shown. (c) IFNγ, IL-13 and IL-17A expression in CD90^hi^, CD90^low^ and CD90^-^ total CD127^+^ ILC and (d) corresponding statistical analyses are outlined. (e) CD90 co-expression with IL-17A or IFNγ in IL-13^+^ ILC and corresponding statistical analyses are shown. (f) Flow cytometry and statistical analysis of CD90 and IL-13 expression in IL-17A^+^ ILC are presented. Data shown are representative of 4 biological replicates.

The abundance of CD90^high^ ILC was greater than that of CD90^-^ and CD90^low^ ILC, but these populations represented ∼30% and 20%, respectively, of the total CD127^+^ ILC population. In order to determine whether CD90^-^ and CD90^low^ ILC were associated with a specific ILC subset, we analysed CD90^-^ and CD90^low^ ILC in *Tbx21*^-/-^ x *Rag2*^-/-^ non-ulcerative colitis (TRnUC) mice. This revealed that the presence of these cells was not dependent on T-bet, and their frequency was not affected. CD90^-^ and CD90^low^ ILC were a relevant source of IFNγ, IL-13 and IL-17A, but still significantly less potent than CD90^high^ ILC in these DSS-treated *Rag2*-deficient mice (Figure 4c, d). DSS-treated TRnUC mice did not have altered frequencies of CD90^-^ and CD90^low^ ILC or IL-13 production in these cells in comparison to DSS-treated *Rag2*^-/-^ mice (Figure 1b-d). However, TRnUC mice had a greater frequency of IL-17A expressing CD90^-^ and CD90^high^ ILC than *Rag2*^-/-^ mice. This could be explained by the far greater cellularity of ILC3 in *Rag2*^-/-^ mice driven by the deficiency of *Tbx21* (Schroeder *et al*., 2021b).

Similar to the observations in DSS-treated WT mice, we detected CD90^-^ and CD90^+^ ILC co-expressing IL-13 and IL-17A (Figure 1c, d, e, f). We also detected more CD90^+^ than CD90^-^ inflammatory IL-13^+^ IL-17A^+^ ILC2 (Figure 1e, f), supporting the notion that inflammatory ILC2 have a CD90^-^ and CD90^+^ phenotype. We also noted that CD90^-^ ILC can also express IL-17A independently of IL-13 (Figure 1f). Interestingly, T-bet-deficiency appears to promote the frequency of CD90^-^ IL-17^+^ among IL-13^+^ ILC2 in these *Rag2*^-/-^ mice.

Functional CD90^-^ and CD90^low^ ILC were also observed in DSS-treated wild-type BALB/c mice (Supplementary Figure 2a, b). In these DSS-treated mice, CD90^high^ ILC were a vastly more significant source of IFNγ, IL-13 and IL-17A in comparison to CD90^-^ and CD90^low^ ILC (Supplementary Figure 2c, d). As observed in Rag2-deficient mice, CD90^-^ and CD90^low^ ILC were able to produce IL-17A and IL-13, but the proportion of CD90^-^ and CD90^low^ ILC producing these cytokines was increased in DSS-treated BALB/c-background *Tbx21*^-/-^ mice (Supplementary Figure 2c, d). These *Tbx21*^-/-^ mice also had an enhanced frequency of IL-17A^+^ CD90^high^ ILC (Supplementary Figure 2c, d). CD90^-^ and CD90^low^ ILC were also detected in DSS-treated wild-type C57BL/6 mice (Supplementary Figure 3a, b). As observed in the other mouse strains, CD90^-^ and CD90^low^ ILC produced IFNγ, IL-13 and IL-17A, although CD90^high^ ILC appeared to be a greater source of these cytokines (Supplementary Figure 3c, d). In contrast to BALB/c background mice, C57BL/6 background *Tbx21*^-/-^ mice did not have a greater prevalence of IL-17A- and IL-13-producing CD90^-^, CD90^low^ or CD90^high^ ILC than WT mice, however, the frequency of CD90^low^ ILC was enhanced significantly (Supplementary Figure 3a, b, c, d).

Furthermore, we did not detect any IFNγ producing IL-13^+^ ILC, but IL-17A was produced among CD90^+^ and CD90^-^ IL-13^+^ ILC2 in DSS-treated BALB/c *Rag2*^-/-^ mice and C57BL/6 WT mice (Figure 1e, f; Supplementary Figure 3e). These data indicate that low expression of CD90 is not a simple marker of inflammatory ILC2 in these mice.

### CD90-negative CD127^+^ ILC have a predominant type 2 phenotype at steady state

Similar to DSS-treated mice (Figure 1, Supplementary Figure 2, and Supplementary Figure 3), most ILC were CD90^high^ in naïve untreated C57BL6 mice. However, CD90^-^ and CD90^low^ ILC populations were also detected in these mice (Supplementary Figure 4a, b). Interestingly, both CD90^-^ and CD90^low^ ILC produced predominately IL-13 and IL-5 and fewer of these cells produced IFNγ and IL-17A (Supplementary Figure 4b-g). Although moderately low, CD90^-^ and CD90^low^ ILC had a significantly greater frequency of IFNγ positivity than CD90^high^ ILC (Supplementary Figure 4c). A similar trend was not observed for IL-17A (Supplementary Figure 4d). IFNγ and IL-17A production was also driven mostly by distinct populations of cells (Supplementary Figure 4b). Further analyses revealed that the prevalence of IL-13^+^ and IL-5^+^ ILC was greater among CD90^high^ and CD90^low^ ILC in comparison to CD90^-^ ILC (Supplementary Figure 4b, e, f, g; Supplementary Figure 5a). *Tbx21*^-/-^ CD90^-^, CD90^low^ and CD90^high^ ILC exhibit greater expression of IL-5 than ILC in WT mice (Supplementary Figure 4b, f, g), which could be explained by one of our previous reports indicating increased cLP ILC2 abundance in *Tbx21*^-/-^ mice (Garrido Mesa *et al*., 2019). Since CD90^-^ and CD90^low^ ILC appeared to be predominately functional ILC2, we sought to determine whether these cells were able to adopt functional characteristics of ILC1 and ILC3. Plasticity of ILC2 allowing expression of T-bet and RORγt is a well-known phenomenon (Bal *et al*., 2020) and inflammatory ILC2 expressing RORγt were reported to have no or lower expression of CD90 in comparison to RORγt-negative natural ILC2 (Huang *et al*., 2015, Flamar *et al*., 2020). We detected minimal co-expression of IL-13 and IL-17A in CD90^-^ and CD90^low^ cLP ILC from naïve WT and *Tbx21*^-/-^ mice indicating the presence of a minor inflammatory ILC2 population (Supplementary Figure 5a). However, we could also find IL-13 and IL-17A co-expressing CD90^high^ ILC. In contrast, virtually no IL-13 and IFNγ co-expressing CD90^-^ and CD90^low^ ILC were detected in these mice (Supplementary Figure 5b).

### CD90 expression in CD127^+^ ILC is controlled by stimulatory cues

Overall, we detected CD90^-^ and CD90^low^ ILC in both untreated and DSS-treated mice. This suggests that CD90 is not a reliable marker for detection of all ILC in the gut. When we analysed CD127 and CD90 co-expression in lineage-negative cLP leukocytes, we noticed that almost all CD90^+^ cLP ILC had a detectable surface expression of CD127 in naïve C57BL/6 WT and DSS-treated C57BL/6 WT, BALB/c WT and BALB/c *Rag2*^-/-^ mice (Figure 2a). For further analyses, KLRG1 was used as a marker of intestinal ILC2 in line with recent publications (Flamar *et al*., 2020, Garrido Mesa *et al*., 2019, Fiancette *et al*., 2021, Schroeder *et al*., 2021b). The use of KLRG1 as a marker for intestinal ILC2 has an advantage over GATA3 as intestinal ILC3 have a low expression of GATA3 and the expression of this transcription factor is variable within the ILC2 population (Zhong *et al*., 2016, Zhong *et al*., 2020, Ricardo-Gonzalez *et al*., 2018). KLRG1^hi^ intestinal ILC as gated in this study require GATA3 for post-developmental maintenance, supporting the notion these cells are ILC2 (Yagi *et al*., 2014). We found that CD90^-^ and CD90^low^ ILC can be detected among both KLRG1^hi^ and KLRG1^-^ cLP ILC, demonstrating that CD90^-^ and CD90^low^ ILC are also components of the non-ILC2 compartment (Figure 2b).

**Figure 2.**
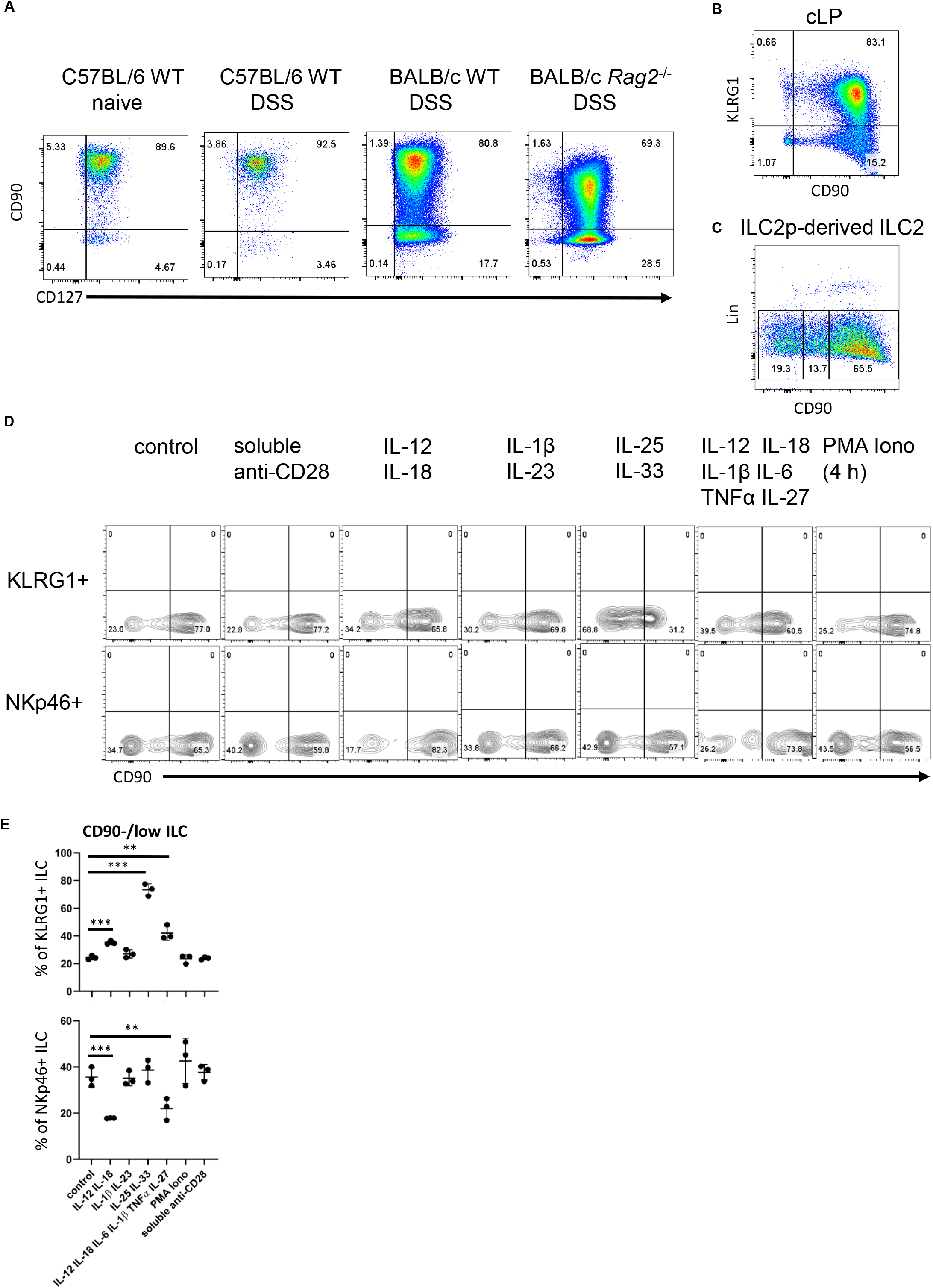
cLP ILC have a variable expression of CD90 depending on stimulatory cues. (a) cLP leukocytes were isolated from untreated C57BL/6 and DSS-treated C57BL/6, BALB/c and Rag-deficient BALB/c mice. CD127 and CD90 co-expression in lineage-negative leukocytes are shown. (b) KLRG1 and CD90 co-expression in cLP CD127^+^ ILC was demonstrated by flow cytometry (n=12). (c) ILC2 were generated from ILC2p stimulated with rIL-7, rSCF and rIL-33, and seeded onto OP9-DL1. CD90^hi^, CD90^low^ and CD90^-^ ILC2 are shown. KLRG1^+^ or KLRG1^-^ CD127^+^ ILC were isolated and stimulated *in vitro* for 48 hours prior to harvest and flow cytometry analyses of KLRG1^+^ or NKp46^+^ ILC, respectively. In addition to a control condition, soluble agonistic anti-CD28 antibodies, IL-12&IL-18, IL-1β&IL23, IL-25&IL-33 or IL-12&IL-18& IL-1β&IL-6&TNFα &IL-27 were used as stimuli. In a separate condition sorted cells were stimulated with PMA and ionomycin in the presence of monensin for the final 4 hours prior to harvesting. (a) Flow cytometry analyses of CD90^hi^ and CD90^low/neg^ CD127^+^ ILC and (b) statistical analyses of CD90^low/neg^ ILC frequencies among KLRG1^+^ or NKp46^+^ cLP ILC are outlined (n=3).

Next, following an established method to develop ILC2 *in vitro*, we seeded bone marrow-derived ILC2 precursors (ILC2p; defined as Lin^-^ CD127^+^ α4β7^hi^ Flt3^-^ CD25^+^) in a 6-day culture on OP9-DL1 stromal cells in the presence of rIL-7, rSCF and rIL-33 (Seehus *et al*., 2016). Strikingly, the Lin^-^ cell population that was generated also exhibited variable levels of CD90 (Figure 2c). Most of the ILC were CD90^high^, but there were also substantial CD90^-^ and CD90^low^ subpopulations.

In order to determine whether CD90 expression can be altered by immunological stimulations, we isolated KLRG1^+^ cLP ILC2 and KLRG1^-^ cLP ILC for *in vitro* culture with OP9-DL1 cells in the presence of distinct cytokines. Strikingly, ILC2 stimulation with IL-25 and IL-33 induced the presence of CD90^-/low^ ILC2 (Figure 2d, e). A similar but less potent effect was observed when IL-12 and IL-18 were added to the culture medium. Additional IL-6/IL-1β/TNFα/IL-27 stimulation did not further alter IL-12/IL-18-mediated CD90^-/low^ ILC2 frequency, while IL-1β/IL-23 stimulation also had no effect. Conversely to ILC2, IL-12/IL-18 stimulation of non-ILC2 in the presence or absence of IL-6/IL-1β/TNFα/IL-27 resulted in fewer CD90^-/low^ NKp46^+^ ILC (Figure 2d, e). IL-1β/IL-23 and IL-25/IL-33 stimulation of these cells had no effect in terms of CD90 expression. Stimulation with PMA and ionomycin or a soluble agonistic anti-CD28 antibody (chosen due to reports of its expression in human ILC (Roan *et al*., 2016, Björklund *et al*., 2016)) also had no effect on the frequency of CD90 expressing ILC2 or NKp46^+^ non-ILC2.

### All ILC subset populations in the intestine exhibit variable levels of CD90

In order to investigate CD90 variation in ILC more closely, we analysed single-cell (sc)RNA-seq data sets from three recent publications: ILC2 from gut, skin, lung, fat and bone marrow (Ricardo-Gonzalez *et al*., (2018)), intestinal ILC2, LTi-like ILC3, NKP46 (NCR)^+^ ILC3, ex-ILC3/ILC1 (Fiancette *et al*., (2021)) and intestinal NK cells, ILC1 and NKp46^+^ ILC3 (Krzywinska *et al*. (2021)) (Figure 3a, b). Visualising clusters of cells that have similar transcriptional profiles using uniform manifold approximation and projection (UMAP) dimensionality reduction and overlaying expression levels of CD90, we found that CD90 expression could be detected across all of the ILC subsets in each dataset (Figure 3c). A pseudotime trajectory analysis of these ILC subsets did not uncover a specific developmental direction from either CD90 high to low expression or vice versa (Figure 3c). Identification of genes up and downregulated in cells positive for CD90 mRNA vs negative/low for CD90 mRNA within each dataset only identified a limited set of genes (Figure 3d). Together with the expression of CD90 across the various cell clusters, this indicates that CD90^-/low^ ILC are not a novel ILC population with their own expression profile. In terms of ILC2, the Fiancette *et al*. data set indicated a specific expression of *Nkg7* in CD90 mRNA-positive cells, but no genes specific for CD90 mRNA-negative/low ILC2 were detected in this data set. In contrast, intestinal CD90 mRNA-negative/low ILC2 exhibited greater expression of *Gzma* (encoding granzyme A) and *Gdd45a, Scin* and *Ctla4*, while intestinal CD90 mRNA-positive ILC2 were characterised by *S100a4, S100a6, Cd3d, Cd3g, Furin* and *Cxcl2* expression. *S100a4* and *S100a6* expression was also detected in CD90 mRNA-positive ILC2 from fat, skin and lung tissue, while *Lgals1* expression was detected in CD90 mRNA-positive ILC2 from lungs, skin and fat tissue. As observed in the Fiancette *et al*. data, *Nkg7* expression as also associated with CD90 mRNA-positive ILC2 from skin and bone marrow, in addition to *Cd7, Ncr1, Klrk1, Ms4a4b* and *Ccl5*. No genes showed consistently higher expression in CD90 mRNA-negative/low cells across all the tissue types but, in the bone marrow, CD90 mRNA-negative/low ILC2 were associated with the expression of *Hbb-bs, Hbb-b7, Hba-a1, Hba-a2* and *S100a8*. The Fiancette *et al*. data set revealed a characteristic expression of *S100a4, S100a6, Pm29* and *Arg1* in CD90 mRNA-positive LTi-like ILC3, while genes specific for CD90 mRNA-negative/low LTi-like ILC3 were not detected. Both the Fiancette *et al*. and Krzywinska *et al*. data sets highlight a specific *Pcp4* expression in CD90 mRNA-positive NKp46^+^ ILC3, while the latter data set also indicate an expression of *Nrgn* in CD90 mRNA-positive NKp46^+^ ILC3 and *Cd74* in CD90^-/low^ NKp46^+^ ILC3. In terms of the ex-ILC3/ILC1 cluster *Tmem176a, Rorc* and *Gda* expression was enhanced in CD90 mRNA-positive cells, while *Ccl5* expression was more abundant in cells in which CD90 mRNA-was absent. In the Krzywinska *et al*. data, CD90 mRNA-positive ILC1 had a characteristic expression of *Il22, Cd83* and *Pxdc1*, while CD90 mRNA-negative/low ILC1 were not defined by specific genes. No genes were found to be upregulated in CD90 mRNA-positive NK cells but *Prf1* and *Gzma* expression was enhanced in CD90 mRNA-negative/low NK cells. Further analyses demonstrated that also only a very few genes were specific for CD90 mRNA-negative/low and CD90 mRNA-high in total ILC and NKp46^+^ ILC (Figure 3e). Similar sets of genes were associated between CD90 mRNA-negative/low and CD90 mRNA-negative total ILC, NKp46^+^ ILC or ILC2 (data not shown) suggesting that these respective populations may be related.

**Figure 3.**
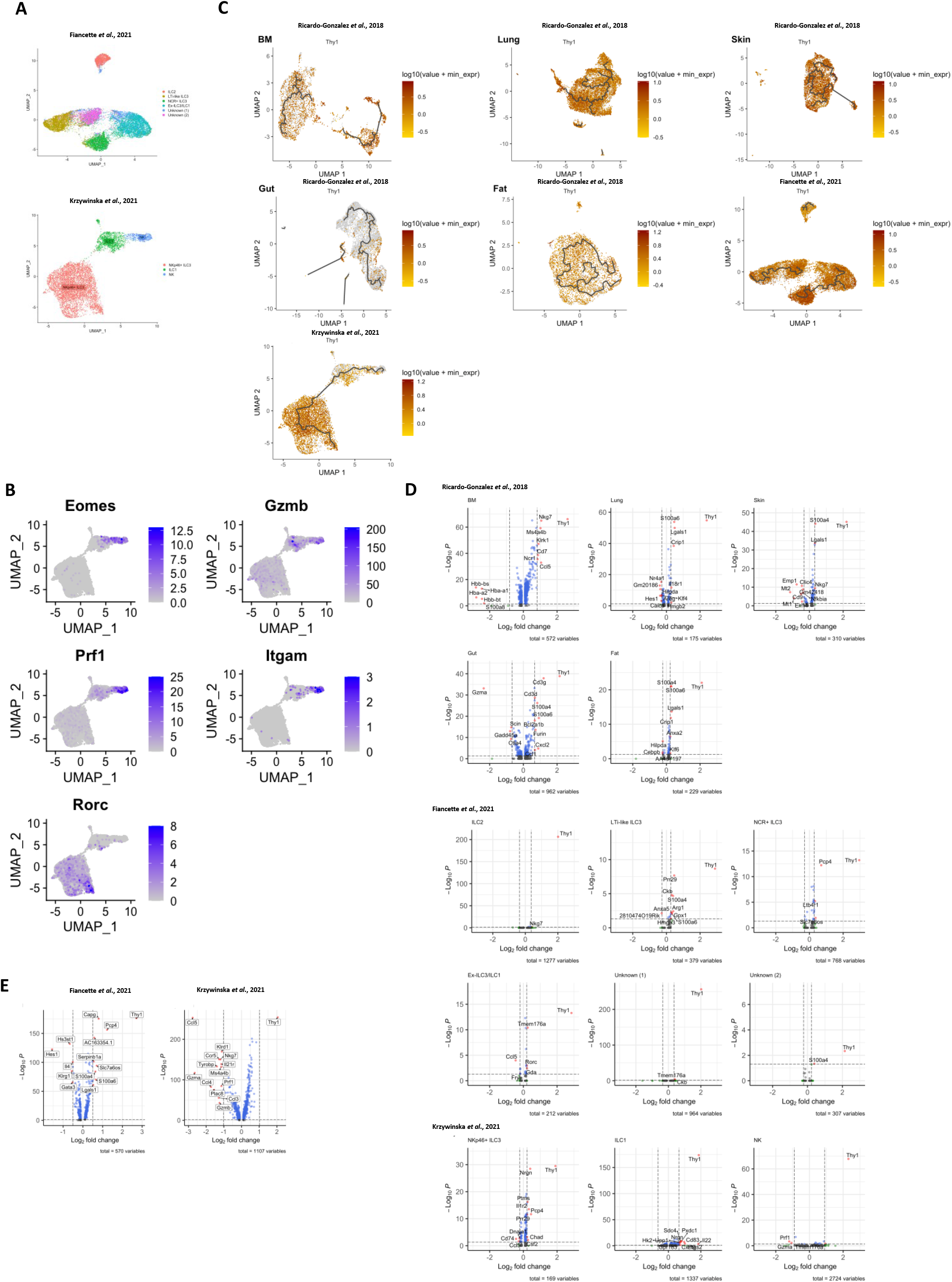
Transcriptome analyses of CD90 expression in intestinal ILC. scRNAseq data sets of intestinal ILC from published studies (Ricardo-Gonzalez *et al*., 2018, Fiancette *et al*., 2021 and Krzywinska *et al*., 2021) were employed to analyse CD90 expression across ILC subsets and its role on the global transcriptional role. (a) UMAP plots of ILC subset annotation in the scRNAseq data sets of the Ricardo-Gonzalez *et al*., (2018), Fiancette *et al*., (2021) and Krzywinska *et al*., (2021) studies. Seurat-based analyses detected distinct ILC2 clusters in the data set published in the Ricardo-Gonzalez *et al*. study. (b) UMAP analyses of gene expression in the ILC subset clusters in the data set obtained from Krzywinska *et al*., (2021). (c) UMAP analysis of CD90 expression intensity in ILC subsets in the respective studies. A trajectory analysis along the CD90 expression intensity was performed in these ILC subsets. (d,e) Vulcano plots of gene expression comparing CD90^high^ ILC versus CD90^low/negative^ ILC subsets as annotated in the respective published data set (d) and total ILC (Fiancette *et al*., (2021)) and total NKp46^+^ ILC (Krzywinska *et al*., (2021)) (e).

### Dysbiosis correlates to ILC1 and ILC3 lymphopenia

Next, we sought to further analyse CD90 expression dynamics in a model of dysbiosis-driven spontaneous colitis in *Tbx21*^-/-^ x *Rag2*^-/-^ ulcerative colitis (TRnUC)mice. We have previously shown that spontaneous colitis in TRUC mice is partially driven by IL-17A-producing CD90^+^ ILC (Powell *et al*., 2012, Powell *et al*., 2015). Hence, it was anticipated that these ILC would also promote inflammation in *Rag2*^-/-^ mice receiving a transfer of feces derived from TRUC mice. These mice developed colitis with decreased body weight and increased colon weight (data not shown). However, in contrast, we detected reduced frequency of DN ILC3, CCR6^+^ ILC3, NKp46^+^ ILC3 and ILC1 (Figure 4a, b). Hence, a ILC2 formed a large proportion of the cLP ILC upon fecal microbial transfer (FMT). In addition to these ILC subset frequency alterations, we detected fewer CD90^high^ and more CD90^low^ cells among the ILC population upon FMT treatment, but the frequency of CD90^-^ ILC was not altered in these mice (Figure 4c, d). Consistent with a greater frequency of ILC2 in FMT-treated mice, cLP ILC production of IL-13 was enhanced, while a significant alteration in the proportion of cells producing IFNγ or IL-17A was not detected (Figure 4e). However, the frequency of IFNγ producing CD90^high^, CD90^low^ and CD90^-^ ILC was much reduced upon the enforced dysbiosis (Figure 4f, g). Furthermore, pathogenic FMT also resulted in a lower frequency of IL-17A^+^ CD90^high^ ILC, while IL-17A production in CD90^low^ and CD90^-^ ILC was not affected. When comparing the proportion of CD90^high^, CD90^low^ and CD90^-^ ILC that produced IFNγ and IL-17A, only a reduction in IFNγ production in CD90^-^ ILC was observed. In contrast to IFNγ and IL-17A, FMT appeared to promote the proportion of IL-13 positive CD90^high^, CD90^low^ and CD90^-^ ILC subsets.

**Figure 4.**
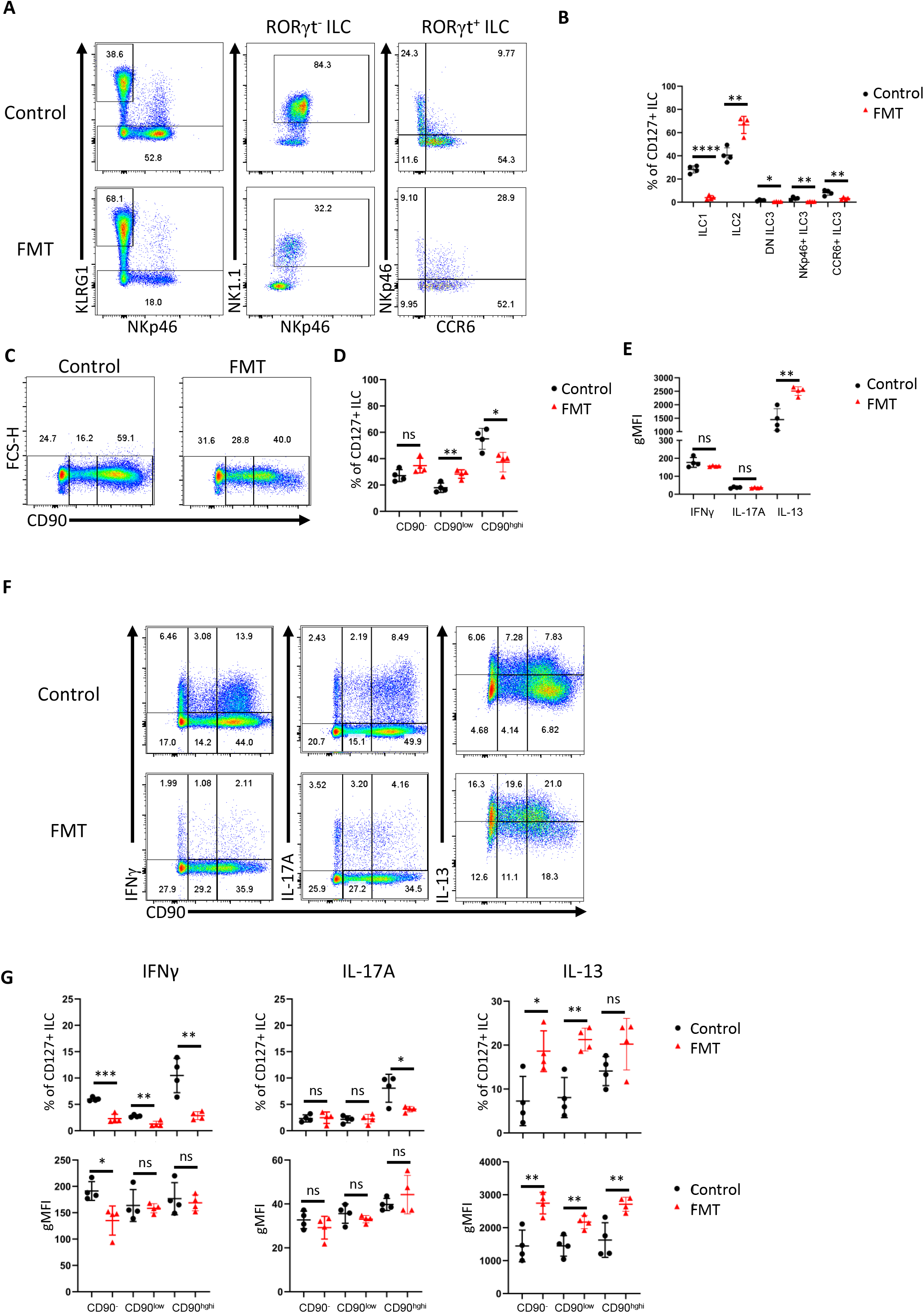
Dysbiosis-triggered appearance of functional cLP ILC with a low expression of CD90. *Rag2*^-/-^ mice were received an FMT using feces derived from TRUC mice and cLP leukocytes were isolated 21 days later from treated and untreated mice. KLRG1^+^ ILC2, KLRG1^-^ RORγt^-^ NKp46^+^ NK1.1^+^ ILC1, KLRG1^-^ RORγt^+^ ILC3 subsets from FMT-treated and untreated control mice were analysed by flow cytometry (A). ILC3 subsets were defined as NKp46^+^ CCR6^-^, CCR6^+^ NKp46^-^ or DN (‘double negative’) in these analyses. A statistical analysis of ILC subset frequency among the whole cLP ILC population is outlined (B). ILC with no or a low or high expression of CD90 were analysed by flow cytometry (C) and a statistical analysis of the frequency of these ILC among the whole ILC population is presented (D). The per cell expression of IFNγ, IL-17A and IL-13 in ILC was analysed statistically (E). IFNγ, IL-17A and IL-13 in CD90^-^, CD90^low^ and CD90^high^ ILC was determined by flow cytometry (F). Related statistical analyses investigating the frequency of respective ILC and the per cell expression of IFNγ, IL-17A and IL-13 in the CD90-, CD90low and CD90high ILC populations is shown (G). Data are representative of 4 biological replicates.

## DISCUSSION

Ever since the discovery of ILC around a decade ago, there have been refinements to the ILC analysis strategy by flow cytometry. This is still an active process, as an increasing number of functional states within the 3 ILC groups are being reported. In the past, many groups have used CD90 as a marker for ILC and CD90-specific antibodies are often employed to deplete ILC *in vivo* (e.g. Powell *et al*., 2012, Martin *et al*., 2017, Rafei-Shamsabadi *et al*., 2018, Castro-Dopico *et al*., 2020, Ray *et al*., 2022, Dobeš *et al*., 2022, Zhou *et al*., 2022). However, our results demonstrate that the use of CD90 to detect and purify ILC has limitations when analysing intestinal populations. In contrast to the notion that CD90 is a pan-ILC marker, the data presented in this study reveal that intestinal ILC can be separated into CD90^-^ and CD90^high^ ILC in addition to CD90^low^ ILC, which are most likely transitional cells. CD90^-^ ILC2 were also detected in the lungs indicating the findings in our study are applicable to ILC from diverse tissues (Loering *et al*., 2021). In our hands, CD127 is a far more reliable marker of ILC than CD90. Virtually all CD90^+^ cLP ILC express CD127, however other reports indicate that pulmonary ILC can lose CD127 *in vivo* and IL-7 downregulates CD127 expression in ILC *in vitro* (Cavagnero *et al*., 2019, Martin *et al*., 2017). Hence, in the absence of better ILC markers, we advise using a combination of CD127 and CD90 to detect ILC.

In BALB/c background mice, CD90^-^ ILC accounted for about a fifth to a third of cLP ILC, and we detected a substantial amount of IFNγ, IL-13 and IL-17A production by these cells upon exposure to DSS- or dysbiosis-elicited colitis. Hence, we believe these findings support the notion that these cells play a relevant role in the ILC response in intestinal tissue. It is out of the scope of this report to define a functionality of CD90 in ILCs, but it was striking to note that whilst ILC2 accounted for most cytokine-producing cLP CD90^-^ ILC in C57BL/6 at steady state, the lack of CD90 expression was not restricted to ILC2. The combination of IL-33 and IL-25, known to activate ILC2, was a potent stimulus for CD90 downregulation in cLP ILC2 *in vitro*, suggesting that low CD90 expression may be an indicator of intestinal ILC2 activity. Interestingly, CD90 expression in pulmonary ILC2 was also shown to drop upon stimulation with IL-33 (Entwistle *et al*., 2020). cLP ILC2 stimulation with IL-12 and IL-18 also enhanced the frequency of CD90^-/low^ cells. It has been reported that IL-12/IL-18 and IL-25/IL-33 can induce ILC2 to express T-bet and RORγt, respectively (Silver *et al*., 2016, Flamar *et al*., 2020). In a model of DSS-induced colitis, we could not associate either IFNγ or IL-17A production by cLP ILC2 with loss of CD90 expression. In contrast to ILC2, CD90 expression was enhanced by IL-12/IL-18-mediated stimulation in NKp46^+^ cLP ILC, which may further indicate that CD90 plays a functional role. Furthermore, in dysbiotic mice we noticed a reduced expression of CD90 in IL-13-producing ILC indicating that CD90 downregulation occurs in activated ILC2 in these mice. Such modified expression of CD90 upon exposure to pathogens is not without precedent. The frequency of intestinal CD90^-^ ILC2 was enhanced in *Hoil1*^-/-^ mice, a mouse model defined by microbe-driven intestinal inflammation (Wood *et al*., 2022). In comparison, an alteration of CD90^+^ ILC2 prevalence was not observed in these mice (Wood *et al*., 2022). Furthermore, *Aspergillus fumigatus*-induced inflammation also leads to the promoted occurrence of pulmonary CD4^+^ T cells with low expression of CD90 (Ulrich *et al*., 2022). In the intestine, variable expression of CD90 can be observed in Vγ7^+^ intraepithelial lymphocytes in addition to conventional CD4^+^ and CD8^+^ T cells (Lei *et al*., 2022, Lo *et al*., 2022).

The functional role of CD90 expression on ILC is unknown and is also ill-defined in other lymphocytes. Known ligands of CD90 are integrins αvβ3, αxβ2, α_M_β2, α_5_β1, α_V_β5, syndecan-4 and CD97, and interactions with binding partners have reported to occur either *in cis* or *in trans* (Wandel *et al*., 2012, Kong *et al*., 2013, Leyton *et al*., 2014, Leyton *et al*., 2019, Pérez *et al*., 2021). *In vitro* studies in in unpolarized and polarized CD4^+^ T cells suggested that CD90 activation with a specific antibody can promote proliferation as well as IFNγ, IL-17A and IL-13 production, in particular in the case of co-stimulation with an agonistic anti-CD28 antibody in the absence of TCR stimulation (Haeryfar *et al*., 2004, Furlong *et al*., 2018). Further work is required to determine the significance of this signalling axis, but, strikingly, scRNA-seq analysis in germinal center (GC) T follicular helper (T_FH_) cells showed distinct transcriptional differences between cells with high expression of CD90 versus cells with low or no expression of CD90 (Yeh *et al*., 2022). These differences included high expression of genes indicative of exocytosis/degranulation in CD90^-/low^ GC T_FH_ cells, and genes relating to chemokine receptors and proliferation in CD90^high^ GC T_FH_ (Yeh *et al*., 2022). Moreover, in addition to CD90^high^ CD8^+^ T cells, splenic CD90^-^ and CD90^low^ CD8^+^ T cells are also a relevant source of IFNγ in a mouse model of LCMV infection (Gather *et al*., 2020). The CD90 extracellular domain has binding sites for αvβ3 and syndecan-4, which may be the basis of a reported *in trans* interaction of CD90 with αvβ3 and syndecan-4 expressed on other cells (Leyton *et al*., 2019). Binding sites for the *in trans* interaction with other integrins or CD97 are yet to be characterised. In addition to *in trans* interactions, αvβ5 is inactive when interacting with CD90 *in cis*, preventing activation of latent TGF-β1 (Zhou *et al*., 2010, Leyton *et al*., 2019). Cis CD90-CD90 interactions have been suggested to promote cluster formation in lipid rafts, which may play a critical for RhoA-dependent signalling, as reported downstream of CD90 (Leyton *et al*., 2019, Pérez *et al*., 2021). Due to its numerous known ligands, CD90 may equip ILC for intercellular interactions with several hematopoietic or non-hematopoietic cell types, but the functional role of CD90 for ILC has still to be defined (Sauzay *et al*., 2019). Interestingly, CD90 was demonstrated to regulate PPARγ expression in adipocytes (Woeller *et al*., 2015), and other groups have reported previously that PPARγ plays an important role in ILC2 functionality (Karagiannis *et al*., 2020, Xiao *et al*., 2021). This study marks the first step to defining CD90 function in ILC by revealing that intestinal ILC can be separated into CD90^+^ and CD90^-^ populations. These data have critical implications for the analysis procedures through which ILC functionality will be uncovered in intestinal tissue.

## METHODS

### Animals

C57BL/6 WT, *Tbx21*^-/-^ (both C57BL/6 and BALB/c background) and *Rag2*^-/-^ (BALB/c background) mice were sourced commercially (Charles River). A colony of colitis-free BALB/c *Rag2*^-/-^ x *Tbx21*^-/-^ (TRnUC) mice was generated from a descendant of the TRUC colony described previously (Garrett *et al*., 2007, Powell *et al*., 2012, Powell *et al*., 2015). All mice were housed in specific pathogen–free facilities at King’s College London Biological Services Unit or at Charles River Laboratories.

### Isolation of cells

cLP leukocytes were isolated using a published method (Gronke *et al*., 2017). Briefly, the epithelium was removed by incubation in HBSS lacking Mg^2+^ or Ca^2+^ (Invitrogen) supplemented with EDTA and HEPES. The tissue was further digested in 2% of foetal calf serum (FCS Gold, PAA Laboratories) supplemented in 0.5 mg/ml collagenase D, 10 μg/ml DNase I and 1.5 mg/ml dispase II (all Roche). The LP lymphocyte-enriched population was harvested from a 40%-80% Percoll (GE Healthcare) gradient.

### Flow Cytometry

Flow cytometry was performed using a standard protocol. Fc receptor blocking was carried out with anti-CD16/32 specific antibodies. A lineage cocktail of antibodies specific for CD3, CD45R, CD19, CD11b, TER-119, Gr-1, CD5 and FcεRI was used for cLP ILC analyses. Live/Dead Fixable Blue Cell Stain Kit (Invitrogen) stain was used to exclude dead cells from the analysis. The cLP ILC gating strategy is outlined in our recent publications (Schroeder *et al*., 2021a, Schroeder *et al*., 2021b). For a complete list of the antibodies used see Table 1. A FoxP3 staining kit (ebioscience) was used for intracellular staining of cytokines and transcription factors. In case of cytokine expression analyses, cells were pre-stimulated with 100 ng/ml PMA and 2 µM ionomycin in the presence of 6 µM monensin for 3-4 hours prior to flow cytometry analysis. Samples were acquired using an LSRFortessa™ cell analyser (Becton Dickinson, USA), and all the data were analysed using FlowJo software (Tree Star, USA).

**Table 1:**
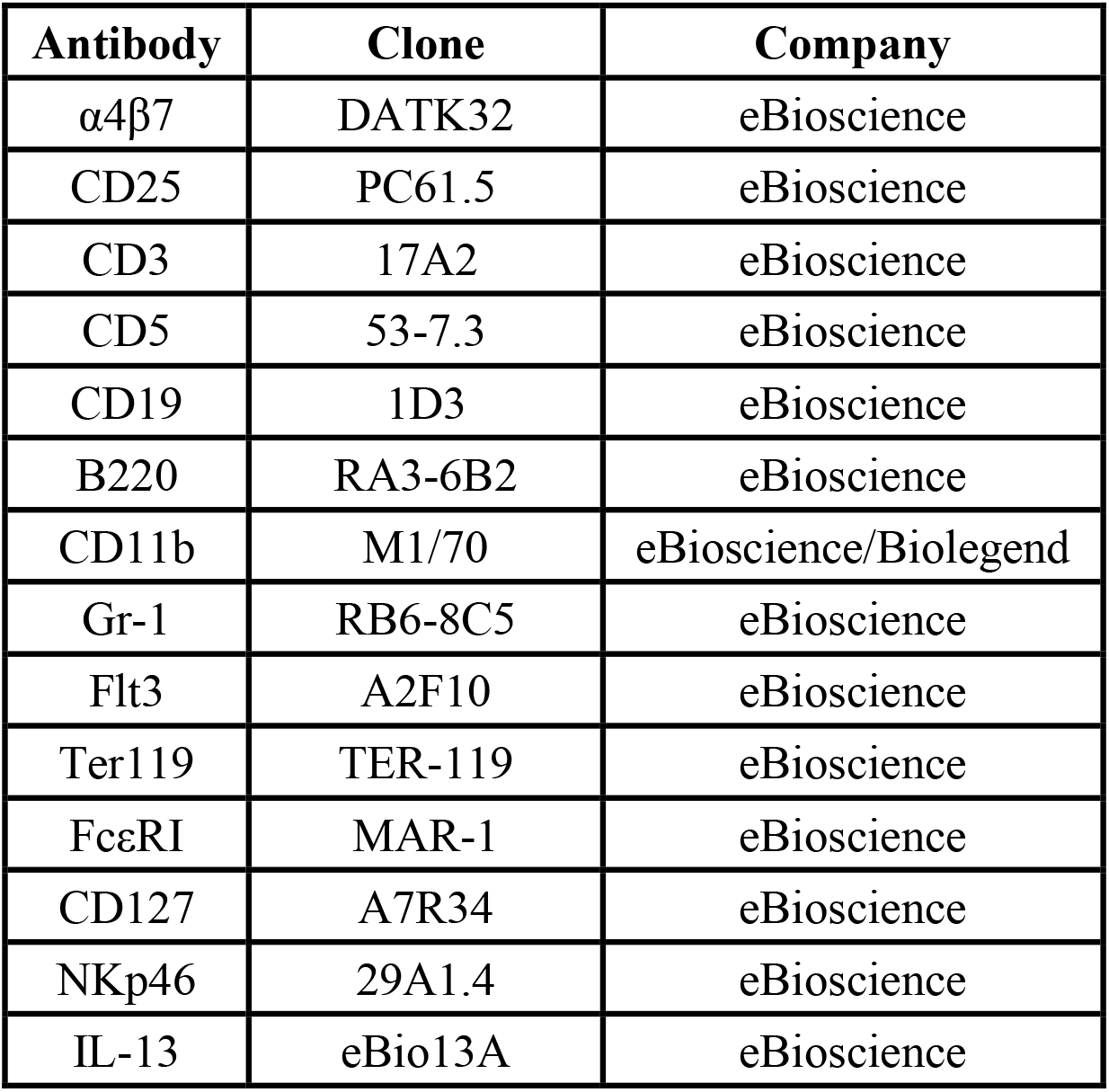
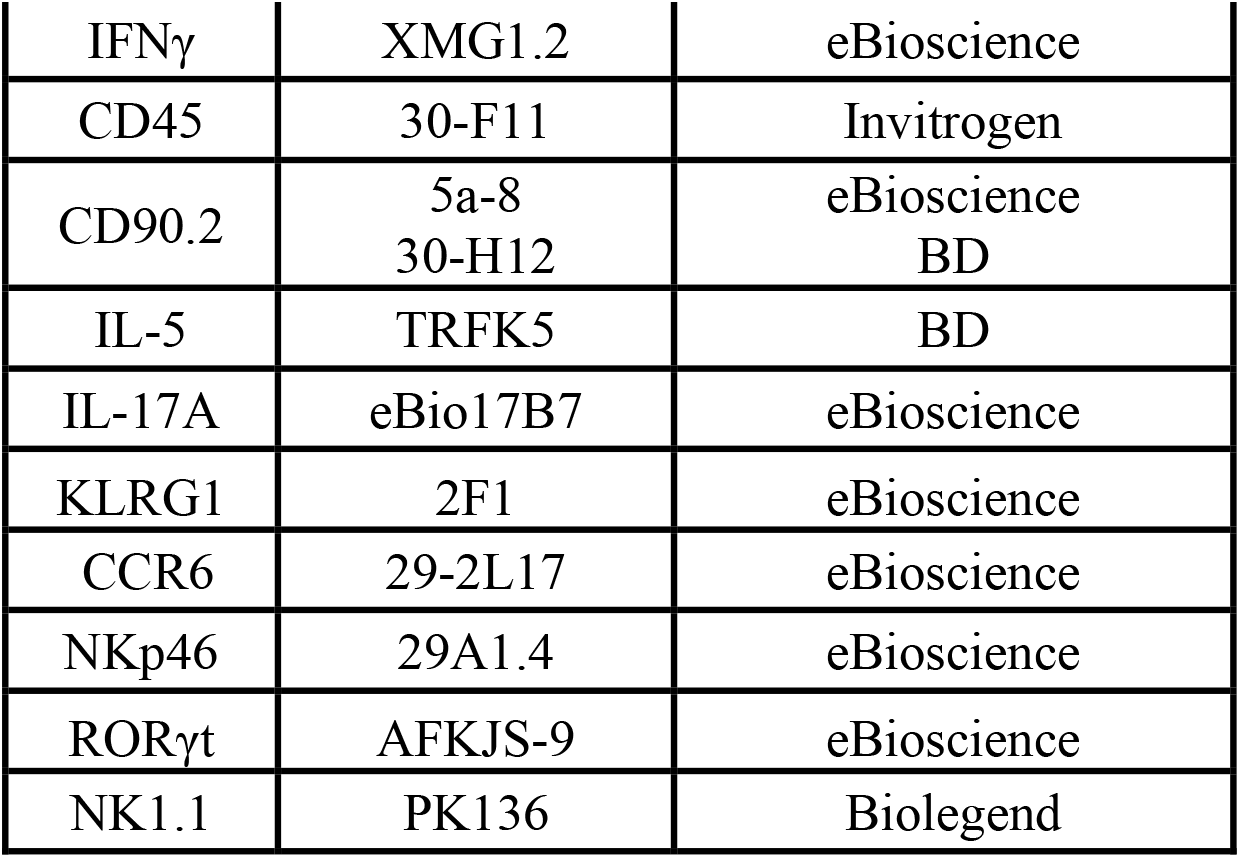
Antibody clones and distributors

| Antibody | Clone | Company |
| --- | --- | --- |
| $\alpha 4\beta 7$ | DATK32 | eBioscience |
| CD25 | PC61.5 | eBioscience |
| CD3 | 17A2 | eBioscience |
| CD5 | 53-7.3 | eBioscience |
| CD19 | 1D3 | eBioscience |
| B220 | RA3-6B2 | eBioscience |
| CD11b | M1/70 | eBioscience/Biolegend |
| Gr-1 | RB6-8C5 | eBioscience |
| Flt3 | A2F10 | eBioscience |
| Ter119 | TER-119 | eBioscience |
| FcεRI | MAR-1 | eBioscience |
| CD127 | A7R34 | eBioscience |
| NKp46 | 29A1.4 | eBioscience |
| IL-13 | eBio13A | eBioscience |
| IFNγ | XMG1.2 | eBioscience |
| CD45 | 30-F11 | Invitrogen |
| CD90.2 | 5a-8<br>30-H12 | eBioscience<br>BD |
| IL-5 | TRFK5 | BD |
| IL-17A | eBio17B7 | eBioscience |
| KLRG1 | 2F1 | eBioscience |
| CCR6 | 29-2L17 | eBioscience |
| NKp46 | 29A1.4 | eBioscience |
| RORγt | AFKJS-9 | eBioscience |
| NK1.1 | PK136 | Biolegend |

### ILC2 generation in OP9-DL1 system

ILC2p were seeded on OP9-DL1 to generate ILC2 using an established method (Seehus *et al*., 2016). Briefly, 7,500 cells were co-cultured with mitomycin pre-treated OP9-DL1 in presence of rmIL-7, rmSCF and rmIL-33 (all 20 ng/ml) for 6 days prior to FACS analysis.

### cLP ILC Sorting and *in vitro* culture

Single-cell suspensions from colonic lamina propria were stained with fluorescently labelled antibodies and isolated using a BD FACSAria III cell sorter (BD Biosciences). Live CD45^+^ Lin^-^ CD127^+^ ILC FACS sorted as KLRG1^+^ and KLRG1^-^ were cultured in DMEM supplemented with 10% FCS, 1xGlutaMax (Gibco), 50 U/ml penicillin, 50 µg/ml streptomycin, 10 mM HEPES, 1x non-essential amino acids (Gibco), 1 mM sodium pyruvate and 50 μM β-mercaptoethanol (Gibco). 20,000 cells were plated per well of a 96-well plate pre-seeded with OP9-DL1 using an established method (Seehus *et al*., 2016, Schroeder *et al*., 2013). The medium was further supplemented with rmIL-7 and rhIL-2 (both at 10 µg/ml) and further recombinant mouse cytokines or anti-CD28 antibody (2µg/ml; clone 37.51) as indicated (all cytokines were used at a final concentration of 10 µg/ml unless indicated otherwise). Cells were harvested and analysed by flow cytometry after 1 day in culture. FACS sort-derived cells from these conditions were harvested and analysed without additional pre-stimulation.

### *In vivo* models

DSS-elicted colitis was induced by adding 3 or 5% DSS (36-50 KDa, MP Biomedicals, Ontario, USA) to the drinking water for 5 days and mice were sacrificed 5-10 days after the beginning of the treatment. To establish a non-colitic control condition, mice were administered sterile drinking water. Regarding all *in vivo* models, body weights and clinical abnormalities were monitored on a daily basis.

For dysbiosis-induced colitis models TRUC mice caecal feces were administered to colitis-free *Rag2*^-/-^ mice via the oral route using a published method and sacrificed 21 days after FMT. (Schroeder *et al*., 2021b). Regarding all *in vivo* models, body weights and clinical abnormalities were monitored on a daily basis.

### Single-cell RNA-seq analysis

Raw expression matrices were obtained from GEO accession GSE117567 (Ricardo-Gonzalez *et al*. 2018) and raw sequencing data were obtained from ArrayExpress accessions E-MTAB-9795 (Fiancette *et al*., 2021) and E-MTAB-11238, (Krzywinska *et al*., 2021). Raw reads were mapped to mm10 using CellRanger 6.0.1. UMAP co-ordinates and clustering metadata was obtained from correspondence with the authors of Fiancette *et al*., 2021 and Krzywinska *et al*., 2021, therefore downstream processing steps can be considered identical to those carried out by the respective authors. For the matrices obtained from Ricardo-Gonzalez *et al*. 2018, cells with over 10% reads mapping to mitochondrial genes and cells with less than 400 genes detected were removed. Each matrix was then normalised using SCTransform (Hafemeister, C. and Satija, R 2019), followed by RunPCA (PCs = 30) and RunUMAP (dims = 30). Shared nearest neighbour and clustering were carried out using FindNeighbours (dims = 30) and FindClusters respectively. NormalizeData was then ran, and this assay was used for downstream visualisation and differential expression analysis using the MAST algorithm (Finak *et al*., 2015). Pseudotime/trajectory analyses were carried out using monocle3 (Trapnell *et al*., 2014 and Qui *et al*., 2017).

### Statistics

Results are expressed as mean ± SEM. Data were analysed using Student’s t-test using GraphPad Prism 5.0 (GraphPad Inc., USA). ns: non-significant; *p < 0.05; **p< 0.01; ***p<0.001; ****p<0.0001.

### Study approval

All animal experiments were performed in accredited facilities in accordance with the UK Animals (Scientific Procedures) Act 1986 (Home Office Licence Numbers PPL: 70/6792, 70/8127 and 70/7869).

## ACKNOWLEDGMENTS

We thank the BRC flow cytometry core team for technical help, and acknowledge financial support from the Department of Health via the NIHR comprehensive Biomedical Research Centre award to Guy’s and St. Thomas’ NHS Foundation Trust in partnership with King’s College London and King’s College Hospital NHS Foundation Trust. We also thank Professor Zúñiga-Pflücker (Sunnybrook Research Institute, University of Toronto) for contributing OP9-DL1 cells. Furthermore, we thank Professor Richard Locksley (University of California San Francisco), Professor Christian Stockmann (University of Zurich), Dr Matthew Hepworth (University of Manchester), Zheng Fan (University of Zurich) and Syed Murtuza Baker (University of Manchester) for providing scRNA-seq data. Work at the CRUK City of London Centre Single Cell Genomics Facility and Cancer Institute Genomics Translational Technology Platform was supported by the CRUK City of London Centre Award [C7893/A26233].

## AUTHOR CONTRIBUTIONS

Study concept and design (JHS, GML, JFN, NP), acquisition of data (JHS, JWL, TZ), bioinformatics analysis (GB), data analysis and interpretation (JHS, GML, JFN, NP, RGJ), obtained funding (GML, NP, JFN), drafting of manuscript (JHS, RGJ), study supervision (GML).

## DISCLOSURE

The authors have no conflict of interest to declare.

## FIGURE LEGENDS

**Supplementary Figure 1:**
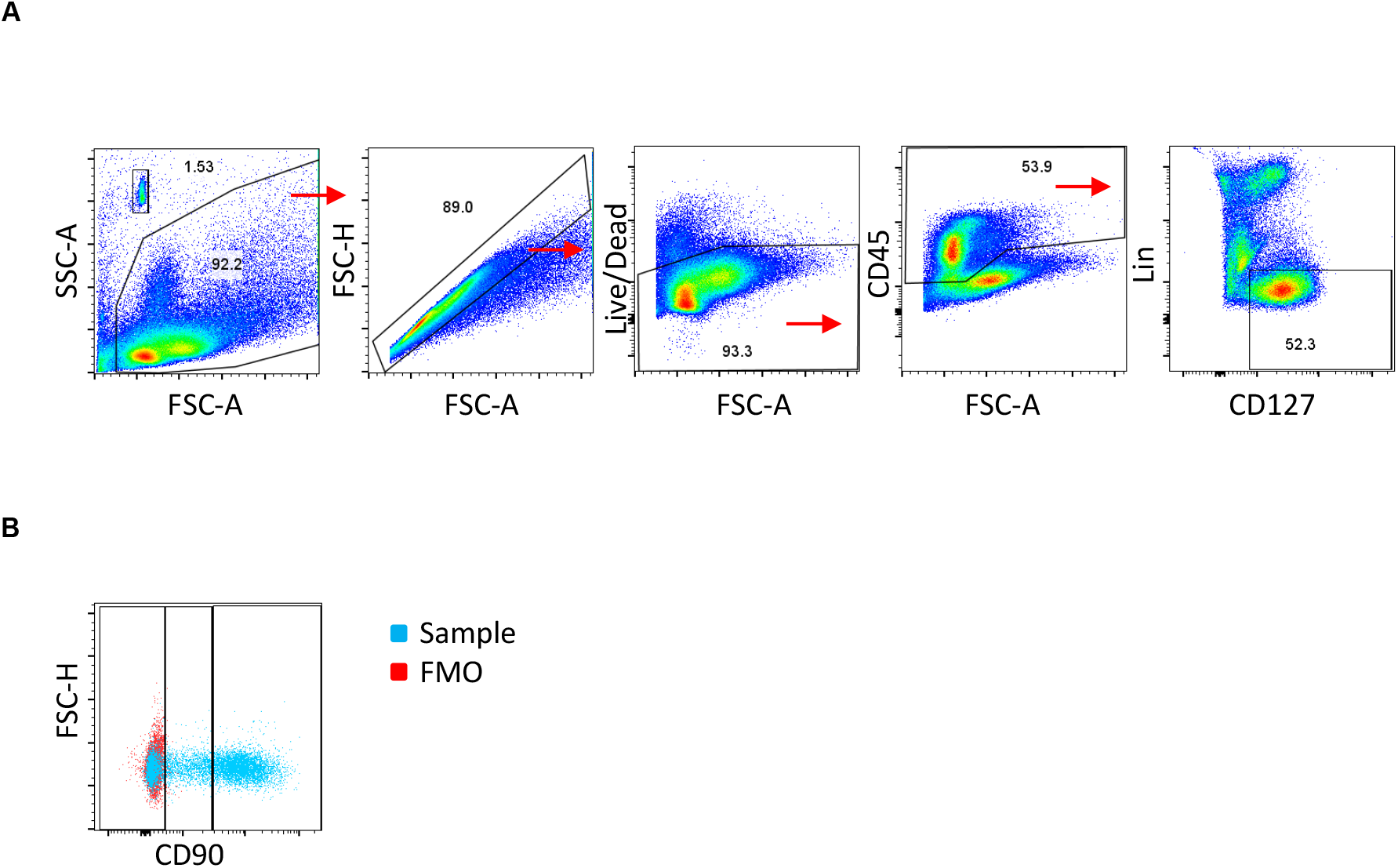
Gating strategy for cLP ILC. Murine cLP ILC were isolated from *Rag2*-deficient mice for flow cytometry analysis. (a) ILC were gated as live single CD45^+^ Lin^-^ CD127^+^ leukocytes. The lineage cocktail contained CD3, CD5, CD19, B220, CD11b, Gr-1, FcεR1 and Ter119. (b) CD90 expression intensity in cLP ILC was evaluated using an FMO control sample.

**Supplementary Figure 2:**
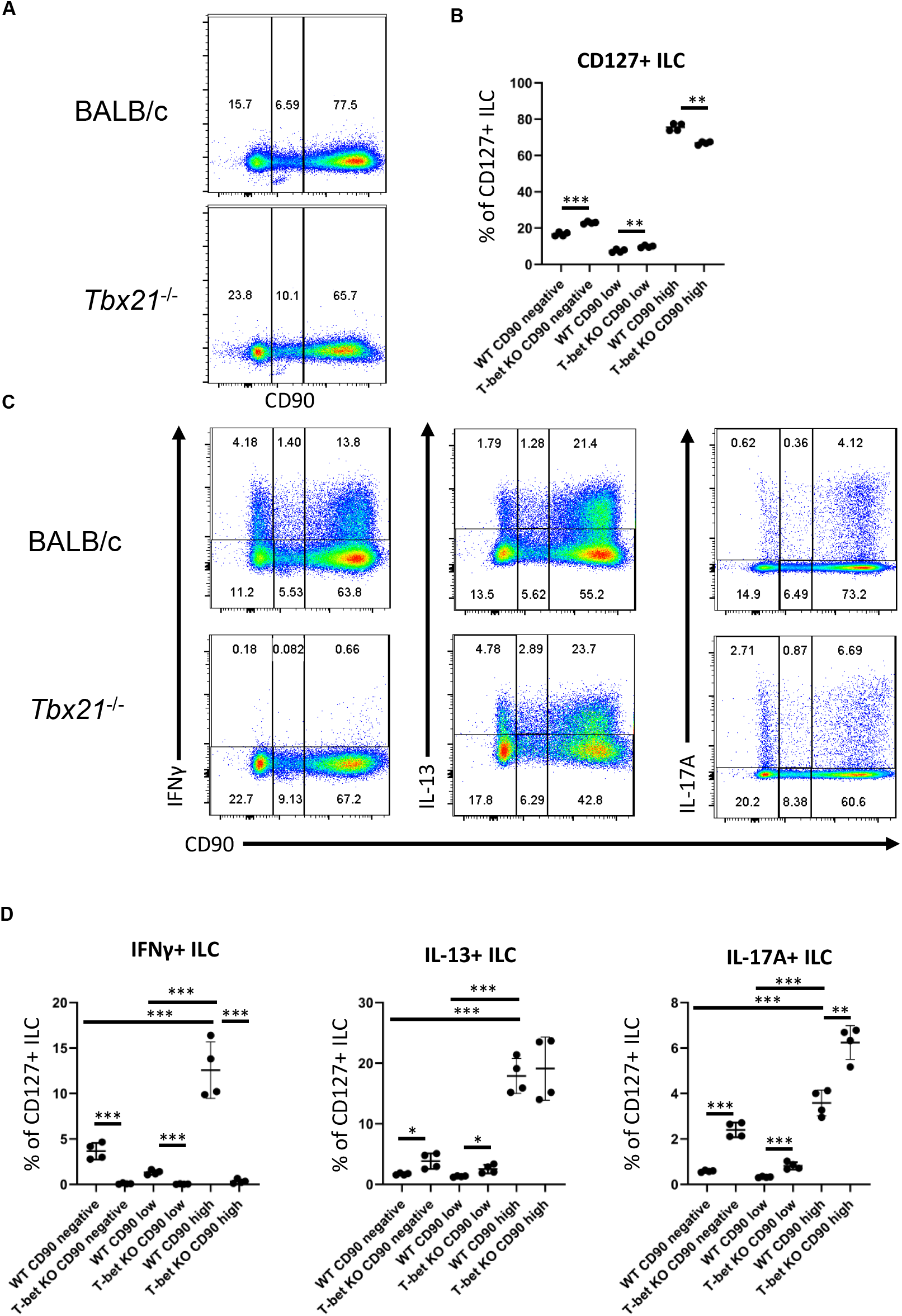
CD90-negative WT cLP CD127^+^ ILC are a source of IFNγ, IL-13 and IL-17A upon DSS treatment of BALB/c mice. cLP ILC from 3% DSS-treated BALB/c WT and *Tbx21*^-/-^ mice were isolated and stimulated with PMA and ionomycin (3 hours) prior to flow cytometry analysis. (a) Frequencies of CD90^hi^, CD90^low^ and CD90^-^ in total CD127^+^ ILC and (b) statistical analyses are outlined. (c) IFNγ, IL-13 and IL-17A expression in CD90^hi^, CD90^low^ and CD90^-^ CD127^+^ ILC and (d) corresponding statistical analyses are shown. Data shown are representative of 4 biological replicates.

**Supplementary Figure 3.**
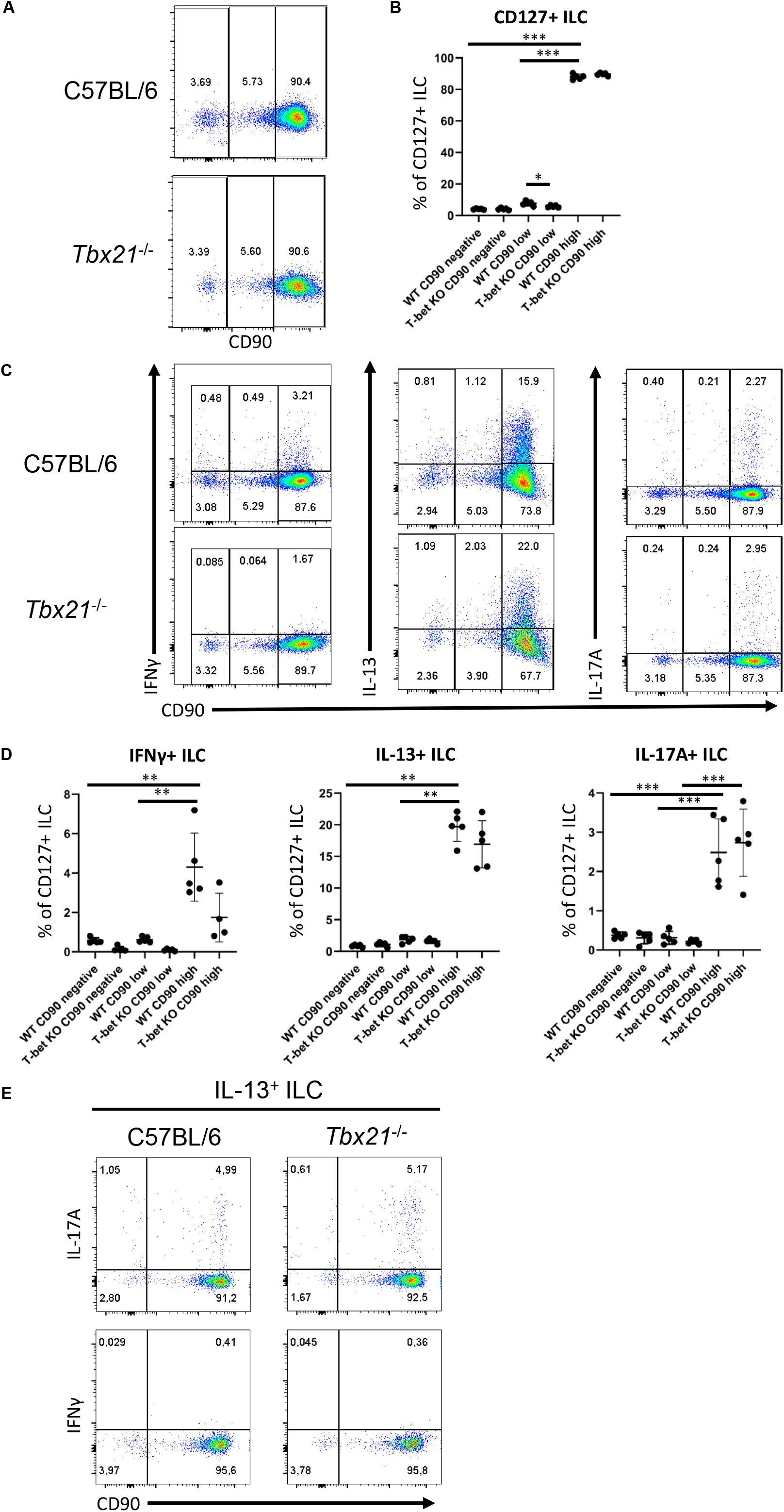
CD90-negative WT cLP CD127^+^ ILC are a minor source of IFNγ, IL-13 and IL-17A during DSS colitis. cLP ILC from 3% DSS-treated C57BL/6 WT and *Tbx21*^-/-^ mice were isolated and stimulated with PMA and ionomycin (3 hours) prior to flow cytometry analysis. (a) Frequencies of CD90^hi^, CD90^low^ and CD90^-^ in total CD127^+^ ILC and (b) statistical analyses are outlined. (c) IFNγ, IL-13 and IL-17A expression in CD90^hi^, CD90^low^ and CD90^-^ CD127^+^ ILC and (d) corresponding statistical analyses are shown. (e) CD90 co-expression with IL-17A or IFNγ in IL-13^+^ ILC is demonstrated. Data shown are representative of 4 biological replicates.

**Supplementary Figure 4.**
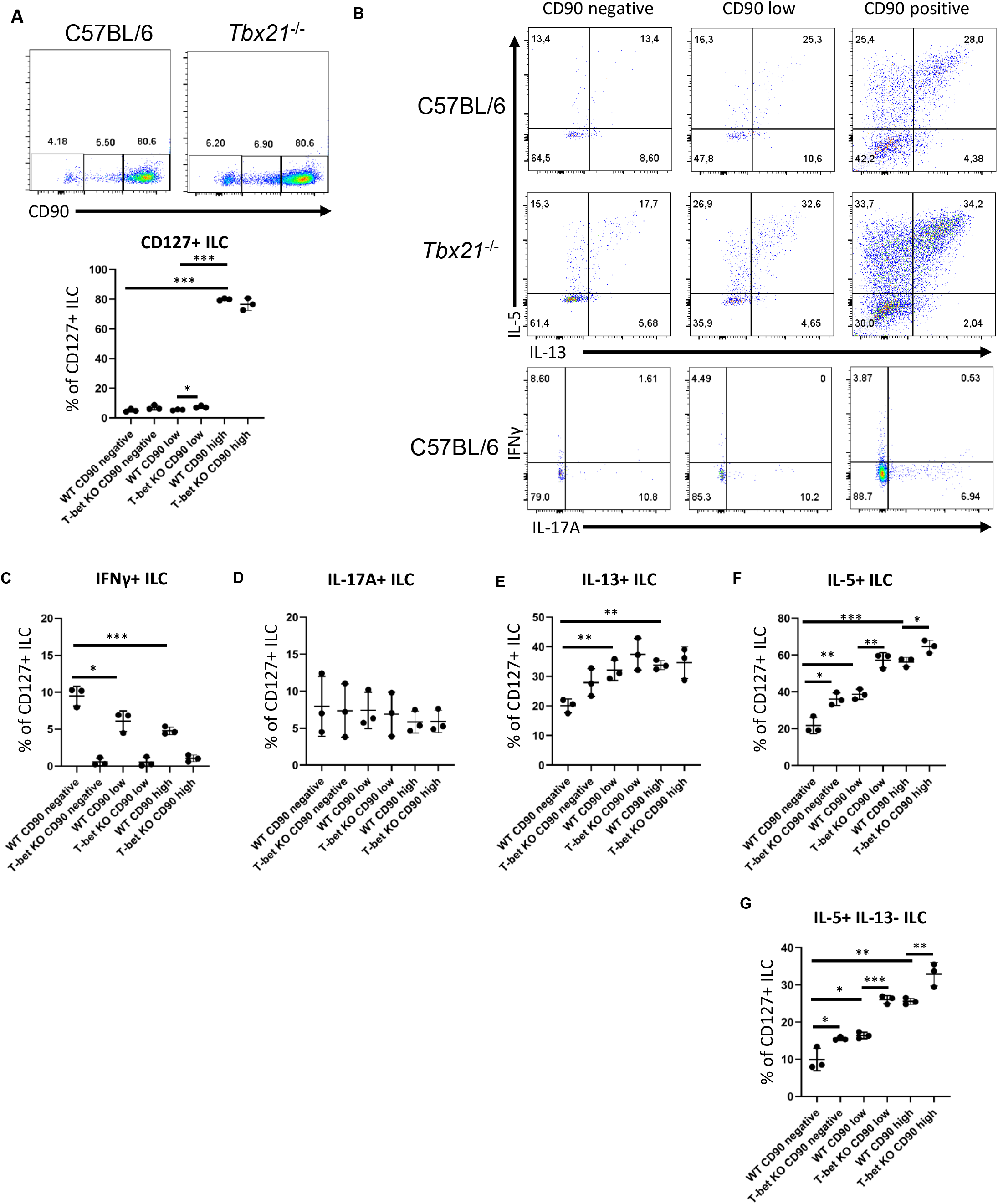
Intestinal CD90-negative CD127^+^ ILC have predominat type 2 phenotype. cLP CD127^+^ ILC were isolated from untreated C57BL/6 WT and *Tbx21*^-/-^ mice and stimulated with PMA and ionomycin (4 hours) prior to flow cytometry analysis. (a) Frequencies of CD90^hi^, CD90^low^ and CD90^-^ in CD127^+^ ILC and statistical analyses are outlined (n=3). (b) IL-13, IL-5, IFNγ and IL-17A expression in CD90^hi^, CD90^low^ and CD90^-^ total CD127^+^ ILC and statistical analyses of (c) IFNγ and (d) IL-17A, (e) IL-13 and (f) IL-5 expression flow cytometry analyses in CD90^hi^, CD90^low^ and CD90^-^ CD127^+^ ILC are illustrated (n=3). (g) Statistical analysis of IL-5^+^ IL-13^-^ CD90^hi^, CD90^low^ and CD90^-^ CD127^+^ ILC (n=3).

**Supplementary Figure 5:**
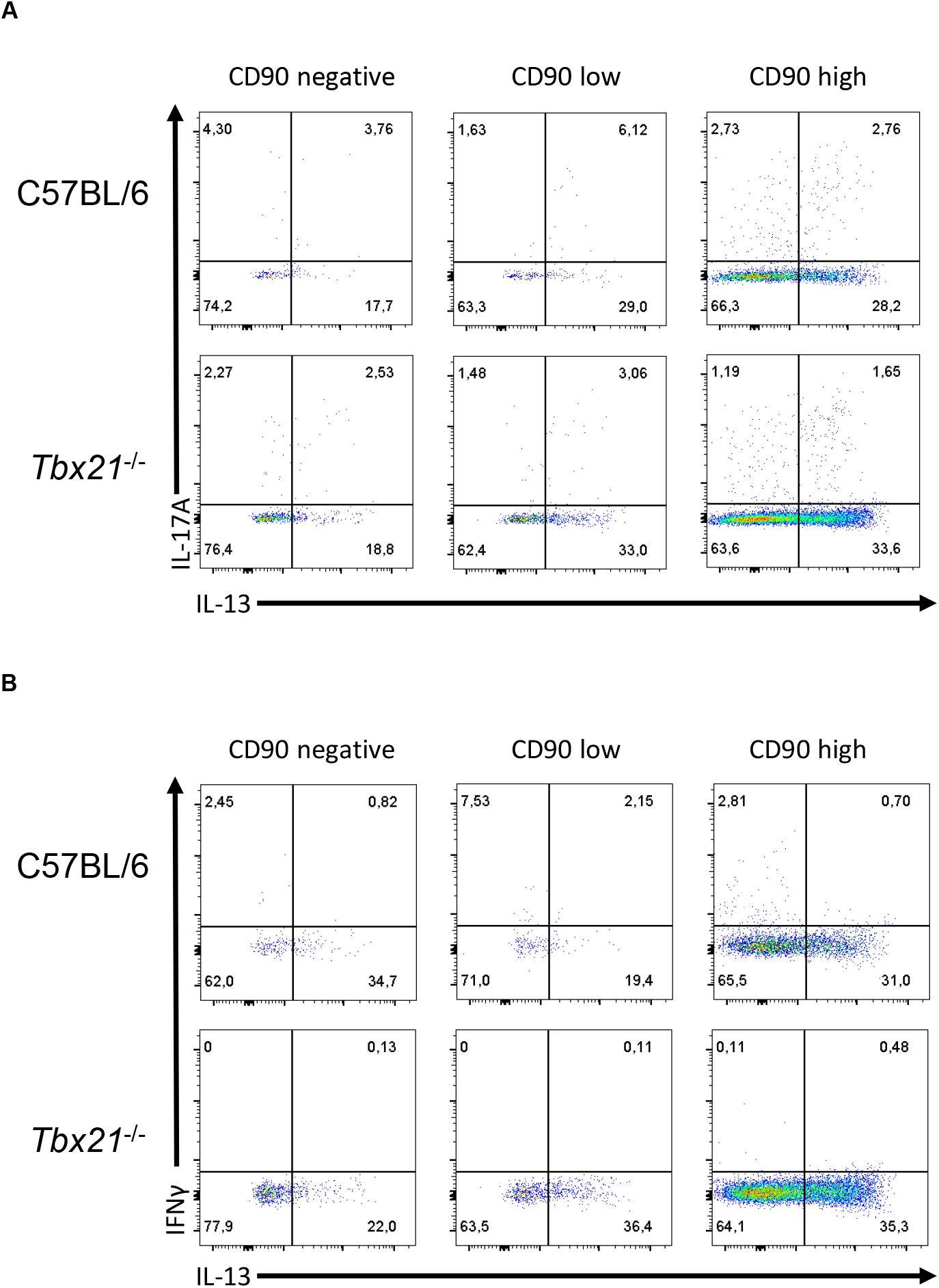
Characterisation of CD90 expression in inflammatory ILC2. cLP CD127^+^ ILC were isolated from untreated C57BL/6 WT and *Tbx21*^-/-^ mice and stimulated with PMA and ionomycin (4 hours) prior to flow cytometry analysis. Flow cytometry analyses of IL-13 co-expression with (a) IL-17A and (b) IFNγ in CD90, CD90^low^ and CD90^-^ CD127^+^ ILC are outlined. Data shown are representative of 3 biological replicates.

## Notes

**Conflict of interest statement** The authors have declared that no conflict of interest exists.

**Financial support:** This study was supported by grants awarded by the Wellcome Trust (091009), the Medical Research Council (MR/M003493/1; MR/K002996/1, all to GML) and RCUK/UKRI Rutherford Fund fellowship (MR/R024812/1) to JFN. Research was also supported by the National Institute for Health Research (NIHR) Biomedical Research Centre at Guy’s and St Thomas and King’s College London. The views expressed are those of the author(s) and not necessarily those of the NHS, the NIHR, or the Department of Health.

### Competing Interest Statement

The authors have declared no competing interest.

### Summary of Updates

v3

## REFERENCES

Alvarez M, Simonetta F, Baker J, Pierini A, Wenokur AS, Morrison AR, Murphy WJ, Negrin RS. (2019) Regulation of murine NK cell exhaustion through the activation of the DNA damage repair pathway. JCI Insight. 5(14):e127729.

Bal SM, Golebski K, Spits H. (2020) Plasticity of innate lymphoid cell subsets. Nat Rev Immunol. 20(9):552–565.

Barman TK, Huber VC, Bonin JL, Califano D, Salmon SL, McKenzie ANJ, Metzger DW. (2022) Viral PB1-F2 and host IFN-γ guide ILC2 and T cell activity during influenza virus infection. Proc Natl Acad Sci U S A. 119(8):e2118535119.

Björklund ÅK, Forkel M, Picelli S, Konya V, Theorell J, Friberg D, Sandberg R, Mjösberg J. (2016) The heterogeneity of human CD127(+) innate lymphoid cells revealed by single-cell RNA sequencing. Nat Immunol. 17(4):451–60.

Bruchard M, Geindreau M, Perrichet A, Truntzer C, Ballot E, Boidot R, Racoeur C, Barsac E, Chalmin F, Hibos C, Baranek T, Paget C, Ryffel B, Rébé C, Paul C, Végran F, Ghiringhelli F. (2022) Recruitment and activation of type 3 innate lymphoid cells promote antitumor immune responses. Nat Immunol. 2022 Feb;23(2):262–274.

Castro-Dopico T, Fleming A, Dennison TW, Ferdinand JR, Harcourt K, Stewart BJ, Cader Z, Tuong ZK, Jing C, Lok LSC, Mathews RJ, Portet A, Kaser A, Clare S, Clatworthy MR. (2020) GM-CSF Calibrates Macrophage Defense and Wound Healing Programs during Intestinal Infection and Inflammation. Cell Rep. 32(1):107857.

Cavagnero KJ, Badrani JH, Naji LH, Amadeo MB, Shah VS, Gasparian S, Pham A, Wang AW, Seumois G, Croft M, Broide DH, Doherty TA. (2019) Unconventional ST2- and CD127-negative lung ILC2 populations are induced by the fungal allergen Alternaria alternata. J Allergy Clin Immunol. 2019 Nov;144(5):1432–1435.e9.

Chen Y, Wang X, Hao X, Li B, Tao W, Zhu S, Qu K, Wei H, Sun R, Peng H, Tian Z. (2022) Ly49E separates liver ILC1s into embryo-derived and postnatal subsets with different functions. J Exp Med. 219(5):e20211805.

Chen W, Chen S, Yan C, Zhang Y, Zhang R, Chen M, Zhong S, Fan W, Zhu S, Zhang D, Lu X, Zhang J, Huang Y, Zhu L, Li X, Lv D, Fu Y, Iv H, Ling Z, Ma L, Jiang H, Long G, Zhu J, Wu D, Wu B, Sun B. (2022) Allergen protease-activated stress granule assembly and gasdermin D fragmentation control interleukin-33 secretion. Nat Immunol. 23(7):1021–1030.

Cox CB, Storm EE, Kapoor VN, Chavarria-Smith J, Lin DL, Wang L, Li Y, Kljavin N, Ota N, Bainbridge TW, Anderson K, Roose-Girma M, Warming S, Arron JR, Turley SJ, de Sauvage FJ, van Lookeren Campagne M. (2021) IL-1R1-dependent signaling coordinates epithelial regeneration in response to intestinal damage. Sci Immunol. 6(59):eabe8856.

Dobeš J, Ben-Nun O, Binyamin A, Stoler-Barak L, Oftedal BE, Goldfarb Y, Kadouri N, Gruper Y, Givony T, Zalayat I, Kováčová K, Böhmová H, Valter E, Shulman Z, Filipp D, Husebye ES, Abramson J. (2022) Extrathymic expression of Aire controls the induction of effective T_H_17 cell-mediated immune response to Candida albicans. Nat Immunol. 23(7):1098–1108.

Entwistle LJ, Gregory LG, Oliver RA, Branchett WJ, Puttur F, Lloyd CM. (2020) Pulmonary Group 2 Innate Lymphoid Cell Phenotype Is Context Specific: Determining the Effect of Strain, Location, and Stimuli. Front Immunol. 10:3114.

Ermann J, Garrett WS, Kuchroo J, Rourida K, Glickman JN, Bleich A, Glimcher LH. (2011) Severity of innate immune-mediated colitis is controlled by the cytokine deficiency-induced colitis susceptibility-1 (Cdcs1) locus. Proc Natl Acad Sci U S A. 108(17):7137–41.

Fachi JL, Pral LP, Dos Santos JAC, Codo AC, de Oliveira S, Felipe JS, Zambom FFF, Câmara NOS, Vieira PMMM, Colonna M, Vinolo MAR. (2021) Hypoxia enhances ILC3 responses through HIF-1α-dependent mechanism. Mucosal Immunol. 14(4):828–841.

Finak G, McDavid A, Yajima M, Deng J, Gersuk V, Shalek AK, Slichter CK, Miller HW, McElrath MJ, Prlic M, Linsley PS, Gottardo R. (2015) MAST: a flexible statistical framework for assessing transcriptional changes and characterizing heterogeneity in single-cell RNA sequencing data. Genome Biol. 16:278.

Flamar AL, Klose CSN, Moeller JB, Mahlakõiv T, Bessman NJ, Zhang W, Moriyama S, Stokic-Trtica V, Rankin LC, Putzel GG, Rodewald HR, He Z, Chen L, Lira SA, Karsenty G, Artis D. (2020) Interleukin-33 Induces the Enzyme Tryptophan Hydroxylase 1 to Promote Inflammatory Group 2 Innate Lymphoid Cell-Mediated Immunity. Immunity. 52(4):606–619.e6.

Furlong S, Coombs MRP, Ghassemi-Rad J, Hoskin DW. (2018) Thy-1 (CD90) Signaling Preferentially Promotes RORγt Expression and a Th17 Response. Front Cell Dev Biol. 6:158.

Garrett WS, Lord GM, Punit S, Lugo-Villarino G, Mazmanian SK, Ito S, Glickman JN, Glimcher LH. (2007) Communicable ulcerative colitis induced by T-bet deficiency in the innate immune system. Cell. 131(1):33–45.

Garrido-Mesa N, Schroeder JH, Stolarczyk E, Gallagher AL, Lo JW, Bailey C, Campbell L, Sexl V, MacDonald TT, Howard JK, Grencis RK, Powell N, Lord GM (2019) T-bet controls intestinal mucosa immune responses via repression of type 2 innate lymphoid cell function. Mucosal Immunol. 12(1):51–63.

Gather R, Aichele P, Goos N, Rohr J, Pircher H, Kögl T, Zeiser R, Hengel H, Schmitt-Gräff A, Weaver C, Ehl S. (2020) Trigger-dependent differences determine therapeutic outcome in murine primary hemophagocytic lymphohistiocytosis. Eur J Immunol. 50(11):1770–1782.

Gillard GO, Bivas-Benita M, Hovav AH, Grandpre LE, Panas MW, Seaman MS, Haynes BF, Letvin NL. (2011) Thy1+ NK cells from vaccinia virus-primed mice confer protection against vaccinia virus challenge in the absence of adaptive lymphocytes. PLoS Pathog. 7(8):e1002141.

Glaubitz J, Wilden A, Golchert J, Homuth G, Völker U, Bröker BM, Thiele T, Lerch MM, Mayerle J, Aghdassi AA, Weiss FU, Sendler M. (2022) In mouse chronic pancreatitis CD25^+^FOXP3^+^ regulatory T cells control pancreatic fibrosis by suppression of the type 2 immune response. Nat Commun. 13(1):4502.

Gronke K, Kofoed-Nielsen M, Diefenbach A. (2017) Isolation and Flow Cytometry Analysis of Innate Lymphoid Cells from the Intestinal Lamina Propria. Methods Mol Biol. 1559:255–265.

Haeryfar SM, Al-Alwan MM, Mader JS, Rowden G, West KA, Hoskin DW. (2003) Thy-1 signaling in the context of costimulation provided by dendritic cells provides signal 1 for T cell proliferation and cytotoxic effector molecule expression, but fails to trigger delivery of the lethal hit. J Immunol. 171(1):69–77.

Hafemeister C, Satija R. (2019) Normalization and variance stabilization of single-cell RNA-seq data using regularized negative binomial regression. Genome Biol. 20(1):296.

Han J, Wan Q, Seo GY, Kim K, El Baghdady S, Lee JH, Kronenberg M, Liu YC. (2022) Hypoxia induces adrenomedullin from lung epithelia, stimulating ILC2 inflammation and immunity. J Exp Med. 219(6):e20211985.

He Y, Xu H, Song C, Koprivsek JJ, Arulanandam B, Yang H, Tao L, Zhong G. (2021) Adoptive Transfer of Group 3-Like Innate Lymphoid Cells Restores Mouse Colon Resistance to Colonization of a Gamma Interferon-Susceptible Chlamydia muridarum Mutant. Infect Immun. 89(2):e00533–20.

He J, Jiang G, Li X, Xiao Q, Chen Y, Xu H, Liu G, Lei A, Zhou P, Shi K, Yang Q, Zhao M, Yao Z, Zhou J. (2022) Bilirubin represents a negative regulator of ILC2 in allergic airway inflammation. Mucosal Immunol. 15(2):314–326.

Huang Y, Guo L, Qiu J, Chen X, Hu-Li J, Siebenlist U, Williamson PR, Urban JF Jr, Paul WE. (2015) IL-25-responsive, lineage-negative KLRG1(hi) cells are multipotential ‘inflammatory’ type 2 innate lymphoid cells. Nat Immunol. 16(2):161–9.

Jowett GM, Read E, Roberts LB, Coman D, Vilà González M, Zabinski T, Niazi U, Reis R, Trieu TJ, Danovi D, Gentleman E, Vallier L, Curtis MA, Lord GM, Neves JF. (2022) Organoids capture tissue-specific innate lymphoid cell development in mice and humans. Cell Rep. 40(9):111281.

Karagiannis F, Masouleh SK, Wunderling K, Surendar J, Schmitt V, Kazakov A, Michla M, Hölzel M, Thiele C, Wilhelm C. (2020) Lipid-Droplet Formation Drives Pathogenic Group 2 Innate Lymphoid Cells in Airway Inflammation. Immunity. 52(4):620–634.e6.

Klose CSN, Artis D. (2020) Innate lymphoid cells control signaling circuits to regulate tissue-specific immunity. Cell Res. 30(6):475–491.

Kong M, Muñoz N, Valdivia A, Alvarez A, Herrera-Molina R, Cárdenas A, Schneider P, Burridge K, Quest AF, Leyton L. (2013) Thy-1-mediated cell-cell contact induces astrocyte migration through the engagement of αVβ3 integrin and syndecan-4. Biochim Biophys Acta. 1833(6):1409–20.

Lei X, Ketelut-Carneiro N, Shmuel-Galia L, Xu W, Wilson R, Vierbuchen T, Chen Y, Reboldi A, Kang J, Edelblum KL, Ward D, Fitzgerald KA. (2022) Epithelial HNF4A shapes the intraepithelial lymphocyte compartment via direct regulation of immune signaling molecules. J Exp Med. 219(8):e20212563.

Leyton L, Hagood JS. (2014) Thy-1 modulates neurological cell-cell and cell-matrix interactions through multiple molecular interactions. Adv Neurobiol. 8:3–20.

Leyton L, Díaz J, Martínez S, Palacios E, Pérez LA, Pérez RD. (2019) Thy-1/CD90 a Bidirectional and Lateral Signaling Scaffold. Front Cell Dev Biol. 7:132.

Liu N, He J, Fan D, Gu Y, Wang J, Li H, Zhu X, Du Y, Tian Y, Liu B, Fan Z. (2022) Circular RNA circTmem241 drives group III innate lymphoid cell differentiation via initiation of Elk3 transcription. Nat Commun. 13(1):4711.

Lo JW, Cozzetto D, Liu Z, Ibraheim H, Sieh J, Olbei M, Alexander J, Miguens Blanco J, Madgwick M, Kudo H, Castro Seoane R, Goldin R, Marchesi R, Korcsmaros T, Lord GM, Powell N. (2022) Immune checkpoint inhibitor-induced colitis is mediated by CXCR6^+^ polyfunctional lymphocytes and is dependent on the IL23/IFNγ axis. Research Square. https://doi.org/10.21203/rs.3.rs-1249584/v1

Loering S, Cameron GJ, Bhatt NP, Belz GT, Foster PS, Hansbro PM, Starkey MR. (2021) Differences in pulmonary group 2 innate lymphoid cells are dependent on mouse age, sex and strain. Immunol Cell Biol. 99(5):542–551.

Martin CE, Spasova DS, Frimpong-Boateng K, Kim HO, Lee M, Kim KS, Surh CD. (2017) Interleukin-7 Availability Is Maintained by a Hematopoietic Cytokine Sink Comprising Innate Lymphoid Cells and T Cells. Immunity. 47(1):171–182.e4.

Mortha A, Chudnovskiy A, Hashimoto D, Bogunovic M, Spencer SP, Belkaid Y, Merad M. (2014) Microbiota-dependent crosstalk between macrophages and ILC3 promotes intestinal homeostasis. Science. 343(6178):1249288.

Peng V, Xing X, Bando JK, Trsan T, Di Luccia B, Collins PL, Li D, Wang WL, Lee HJ, Oltz EM, Wang T, Colonna M. (2022) Whole-genome profiling of DNA methylation and hydroxymethylation identifies distinct regulatory programs among innate lymphocytes. Nat Immunol. 23(4):619–631.

Peng V, Cao S, Trsan T, Bando JK, Avila-Pacheco J, Cleveland JL, Clish C, Xavier RJ, Colonna M. (2022) Ornithine decarboxylase supports ILC3 responses in infectious and autoimmune colitis through positive regulation of IL-22 transcription. Proc Natl Acad Sci U S A. 119(45):e2214900119.

Potempa M, Aguilar OA, Gonzalez-Hinojosa MDR, Tenvooren I, Marquez DM, Spitzer MH, Lanier LL. (2022) Influence of Self-MHC Class I Recognition on the Dynamics of NK Cell Responses to Cytomegalovirus Infection. J Immunol. 208(7):1742–1754.

Powell N, Walker AW, Stolarczyk E, Canavan JB, Gökmen MR, Marks E, Jackson I, Hashim A, Curtis MA, Jenner RG, Howard JK, Parkhill J, MacDonald TT, Lord GM. (2012) The transcription factor T-bet regulates intestinal inflammation mediated by interleukin-7 receptor+ innate lymphoid cells. Immunity. 37(4):674–84.

Powell N, Lo JW, Biancheri P, Vossenkämper A, Pantazi E, Walker AW, Stolarczyk E, Ammoscato F, Goldberg R, Scott P, Canavan JB, Perucha E, Garrido-Mesa N, Irving PM, Sanderson JD, Hayee B, Howard JK, Parkhill J, MacDonald TT, Lord GM. (2015) Interleukin 6 Increases Production of Cytokines by Colonic Innate Lymphoid Cells in Mice and Patients With Chronic Intestinal Inflammation. Gastroenterology.149(2):456–67.e15.

Qiu X, Mao Q, Tang Y, Wang L, Chawla R, Pliner HA, Trapnell C. (2017) Reversed graph embedding resolves complex single-cell trajectories. Nat Methods. 14(10):979–982.

Rafei-Shamsabadi DA, van de Poel S, Dorn B, Kunz S, Martin SF, Klose CSN, Arnold SJ, Tanriver Y, Ebert K, Diefenbach A, Halim TYF, McKenzie ANJ, Jakob T. (2018) Lack of Type 2 Innate Lymphoid Cells Promotes a Type I-Driven Enhanced Immune Response in Contact Hypersensitivity. J Invest Dermatol. 138(9):1962–1972.

Ray JL, Shaw PK, Postma B, Beamer CA, Holian A. (2022) Nanoparticle-Induced Airway Eosinophilia Is Independent of ILC2 Signaling but Associated With Sex Differences in Macrophage Phenotype Development. J Immunol. 208(1):110–120.

Ricardo-Gonzalez RR, Van Dyken SJ, Schneider C, Lee J, Nussbaum JC, Liang HE, Vaka D, Eckalbar WL, Molofsky AB, Erle DJ, Locksley RM. (2018) Tissue signals imprint ILC2 identity with anticipatory function. Nat Immunol. 19(10):1093–1099.

Riding AM, Loudon KW, Guo A, Ferdinand JR, Lok LSC, Richoz N, Stewart A, Castro-Dopico T, Tuong ZK, Fiancette R, Bowyer GS, Fleming A, Gillman ES, Suchanek O, Mahbubani KT, Saeb-Parsy K, Withers D, Dougan G, Clare S, Clatworthy MR. (2022) Group 3 innate lymphocytes make a distinct contribution to type 17 immunity in bladder defence. iScience. 25(7):104660.

Roan F, Stoklasek TA, Whalen E, Molitor JA, Bluestone JA, Buckner JH, Ziegler SF. (2016) CD4+ Group 1 Innate Lymphoid Cells (ILC) Form a Functionally Distinct ILC Subset That Is Increased in Systemic Sclerosis. J Immunol. 196(5):2051–2062.

Roberts LB, Schnoeller C, Berkachy R, Darby M, Pillaye J, Oudhoff MJ, Parmar N, Mackowiak C, Sedda D, Quesniaux V, Ryffel B, Vaux R, Gounaris K, Berrard S, Withers DR, Horsnell WGC, Selkirk ME. (2021) Acetylcholine production by group 2 innate lymphoid cells promotes mucosal immunity to helminths. Sci Immunol. 6(57):eabd0359.

Salimi M, Wang R, Yao X, Li X, Wang X, Hu Y, Chang X, Fan P, Dong T, Ogg G. (2018) Activated innate lymphoid cell populations accumulate in human tumour tissues. BMC Cancer. 18(1):341.

Sauzay C, Voutetakis K, Chatziioannou A, Chevet E, Avril T. (2019) CD90/Thy-1, a Cancer-Associated Cell Surface Signaling Molecule. Front Cell Dev Biol. 7:66.

Schmalzl A, Leupold T, Kreiss L, Waldner M, Schürmann S, Neurath MF, Becker C, Wirtz S. (2022) Interferon regulatory factor 1 (IRF-1) promotes intestinal group 3 innate lymphoid responses during Citrobacter rodentium infection. Nat Commun. 13(1):5730.

Schroeder JH, Bell LS, Janas ML, Turner M. (2013) Pharmacological inhibition of glycogen synthase kinase 3 regulates T cell development in vitro. PLoS One. 8(3):e58501

Schroeder JH, Roberts LB, Meissl K, Lo JW, Hromadová D, Hayes K,, Zabinski T, Read E, Moreira Heliodoro C, Reis R, Neves J F, Howard JK, Strobl B, Grencis RK, Lord GM. (2021a) Sustained post-developmental T-bet expression is critical for the maintenance of type one innate lymphoid cells in vivo. Front Immunol. 12:760198.

Schroeder JH, Meissl K, Hromadová D, Lo JW, Neves J F, Helmby H, Powell N, Strobl B, Lord GM (2021b) T-bet controls cellularity of intestinal group 3 innate lymphoid cells. Front Immunol. 11:623324.

Schroeder JH, Howard JK, Lord GM. (2022) Transcription factor-driven regulation of ILC1 and ILC3. Trends Immunol. 43(7):564–579.

Schwartz C, Khan AR, Floudas A, Saunders SP, Hams E, Rodewald HR, McKenzie ANJ, Fallon PG. (2017) ILC2s regulate adaptive Th2 cell functions via PD-L1 checkpoint control. J Exp Med. 214(9):2507–2521.

Seehus CR, Aliahmad P, de la Torre B, Iliev ID, Spurka L, Funari VA, Kaye (2015) The development of innate lymphoid cells requires TOX-dependent generation of a common innate lymphoid cell progenitor. J.Nat Immunol. 16(6):599–608.

Seehus C, Kaye J. (2016) In vitro Differentiation of Murine Innate Lymphoid Cells from Common Lymphoid Progenitor Cells. Bio Protoc. 6(6):e1770.

Sheikh A, Lu J, Melese E, Seo JH, Abraham N. (2022) IL-7 induces type 2 cytokine response in lung ILC2s and regulates GATA3 and CD25 expression. J Leukoc Biol. 112(5):1105–1113.

Silver JS, Kearley J, Copenhaver AM, Sanden C, Mori M, Yu L, Pritchard GH, Berlin AA, Hunter CA, Bowler R, Erjefalt JS, Kolbeck R, Humbles AA. (2016) Inflammatory triggers associated with exacerbations of COPD orchestrate plasticity of group 2 innate lymphoid cells in the lungs. Nat Immunol. 17(6):626–35.

Sparano C, Solís-Sayago D, Vijaykumar A, Rickenbach C, Vermeer M, Ingelfinger F, Litscher G, Fonseca A, Mussak C, Mayoux M, Friedrich C, Nombela-Arrieta C, Gasteiger G, Becher B, Tugues S. (2022) Embryonic and neonatal waves generate distinct populations of hepatic ILC1s. Sci Immunol. 7(75):eabo6641.

Suntharalingam G, Perry MR, Ward S, Brett SJ, Castello-Cortes A, Brunner MD, Panoskaltsis N. (2006) Cytokine storm in a phase 1 trial of the anti-CD28 monoclonal antibody TGN1412. N Engl J Med. 355(10):1018–28.

Trapnell C, Cacchiarelli D, Grimsby J, Pokharel P, Li S, Morse M, Lennon NJ, Livak KJ, Mikkelsen TS, Rinn JL. (2014) The dynamics and regulators of cell fate decisions are revealed by pseudotemporal ordering of single cells. Nat Biotechnol. 32(4):381–386.

Ualiyeva S, Lemire E, Aviles EC, Wong C, Boyd AA, Lai J, Liu T, Matsumoto I, Barrett NA, Boyce JA, Haber AL, Bankova LG. (2021) Tuft cell-produced cysteinyl leukotrienes and IL-25 synergistically initiate lung type 2 inflammation. Sci Immunol. 6(66):eabj0474.

Ulrich BJ, Kharwadkar R, Chu M, Pajulas A, Muralidharan C, Koh B, Fu Y, Gao H, Hayes TA, Zhou HM, Goplen NP, Nelson AS, Liu Y, Linnemann AK, Turner MJ, Licona-Limón P, Flavell RA, Sun J, Kaplan MH. (2022) Allergic airway recall responses require IL-9 from resident memory CD4^+^ T cells. Sci Immunol. 7(69):eabg9296.

Wallrapp A, Riesenfeld SJ, Burkett PR, Abdulnour RE, Nyman J, Dionne D, Hofree M, Cuoco MS, Rodman C, Farouq D, Haas BJ, Tickle TL, Trombetta JJ, Baral P, Klose CSN, Mahlakõiv T, Artis D, Rozenblatt-Rosen O, Chiu IM, Levy BD, Kowalczyk MS, Regev A, Kuchroo VK. (2017) The neuropeptide NMU amplifies ILC2-driven allergic lung inflammation. Nature. 549(7672):351–356.

Wandel E, Saalbach A, Sittig D, Gebhardt C, Aust G. (2012) Thy-1 (CD90) is an interacting partner for CD97 on activated endothelial cells. J Immunol. 188(3):1442–50.

Woeller CF, O’Loughlin CW, Pollock SJ, Thatcher TH, Feldon SE, Phipps RP. (2015) Thy1 (CD90) controls adipogenesis by regulating activity of the Src family kinase, Fyn. FASEB J. 29(3):920–31.

Wood MJ, Marshall JN, Hartley VL, Liu TC, Iwai K, Stappenbeck TS, MacDuff DA. (2022) HOIL1 regulates group 2 innate lymphoid cell numbers and type 2 inflammation in the small intestine. Mucosal Immunol. 15(4):642–655.

Wu X, Kasmani MY, Zheng S, Khatun A, Chen Y, Winkler W, Zander R, Burns R, Taparowsky EJ, Sun J, Cui W. (2022a) BATF promotes group 2 innate lymphoid cell-mediated lung tissue protection during acute respiratory virus infection. Sci Immunol. 7(67):eabc9934.

Wu X, Khatun A, Kasmani MY, Chen Y, Zheng S, Atkinson S, Nguyen C, Burns R, Taparowsky EJ, Salzman NH, Hand TW, Cui W. (2022b) Group 3 innate lymphoid cells require BATF to regulate gut homeostasis in mice. J Exp Med. 219(11):e20211861.

Xiao Q, He J, Lei A, Xu H, Zhang L, Zhou P, Jiang G, Zhou J. (2021) PPARγ enhances ILC2 function during allergic airway inflammation via transcription regulation of ST2. Mucosal Immunol. 14(2):468–478.

Xiao Q, Han X, Liu G, Zhou D, Zhang L, He J, Xu H, Zhou P, Yang Q, Chen J, Zhou J, Jiang G, Yao Z. (2022) Adenosine restrains ILC2-driven allergic airway inflammation vi A2A receptor. Mucosal Immunol. 15(2):338–350.

Yagi R, Zhong C, Northrup DL, Yu F, Bouladoux N, Spencer S, Hu G, Barron L, Sharma S, Nakayama T, Belkaid Y, Zhao K, Zhu J. (2014) The transcription factor GATA3 is critical for the development of all IL-7Rα-expressing innate lymphoid cells. Immunity. 40(3):378–88.

Yeh CH, Finney J, Okada T, Kurosaki T, Kelsoe G. (2022) Primary germinal center-resident T follicular helper cells are a physiologically distinct subset of CXCR5^hi^PD-1^hi^ T follicular helper cells. Immunity. 55(2):272–289.e7.

Zhong C, Cui K, Wilhelm C, Hu G, Mao K, Belkaid Y, Zhao K, Zhu J. (2016) Group 3 innate lymphoid cells continuously require the transcription factor GATA-3 after commitment. Nat Immunol. 17(2):169–78.

Zhong C, Zheng M, Cui K, Martins AJ, Hu G, Li D, Tessarollo L, Kozlov S, Keller JR, Tsang JS, Zhao K, Zhu J. (2020) Differential Expression of the Transcription Factor GATA3 Specifies Lineage and Functions of Innate Lymphoid Cells. Immunity. 52(1):83–95.e4.

Zhou Y, Hagood JS, Lu B, Merryman WD, Murphy-Ullrich JE. (2010) Thy-1-integrin alphav beta5 interactions inhibit lung fibroblast contraction-induced latent transforming growth factor-beta1 activation and myofibroblast differentiation. J Biol Chem. 285(29):22382–93.

Zhou L, Zhou W, Joseph AM, Chu C, Putzel GG, Fang B, Teng F, Lyu M, Yano H, Andreasson KI, Mekada E, Eberl G, Sonnenberg GF. (2022) Group 3 innate lymphoid cells produce the growth factor HB-EGF to protect the intestine from TNF-mediated inflammation. Nat Immunol. 23(2):251–261.

